# Synaptic co-transmission of acetylcholine and GABA regulates hippocampal states

**DOI:** 10.1101/193318

**Authors:** Virág T. Takács, Csaba Cserép, Dániel Schlingloff, Balázs Pósfai, András Szőnyi, Katalin E. Sos, Zsuzsanna Környei, Ádám Dénes, Attila I. Gulyás, Tamás F. Freund, Gábor Nyiri

**Affiliations:** Laboratory of Cerebral Cortex Research, Institute of Experimental Medicine, Hungarian Academy of Sciences, Budapest, H-1083, Hungary.; *Momentum* Laboratory of Neuroimmunology, Institute of Experimental Medicine, Hungarian Academy of Sciences, Budapest, H-1083, Hungary.; János Szentágothai Doctoral School of Neurosciences, Semmelweis University, Budapest, H-1085, Hungary.

## Abstract

The basal forebrain cholinergic system is widely assumed to control cortical functions via non-synaptic transmission of a single neurotransmitter, acetylcholine. Yet, using immune-electron tomographic, molecular anatomical, optogenetic and physiological techniques, we find that mouse hippocampal cholinergic terminals invariably establish synapses and their vesicles dock at synapses only. We demonstrate that these synapses do not co-release but co-transmit GABA and acetylcholine via different vesicles, whose release is triggered by distinct calcium channels. This co-transmission evokes fast composite postsynaptic potentials, which are mutually cross-regulated by presynaptic auto-receptors and display different short-term plasticity. The GABAergic component alone effectively suppresses hippocampal sharp wave-ripples and epileptiform activity. The synaptic nature of the forebrain cholinergic system with differentially regulated, fast, GABAergic and cholinergic co-transmission suggests a hitherto unrecognized level of synaptic control over cortical states. This novel model of hippocampal cholinergic neurotransmission could form the basis for alternative pharmacotherapies after cholinergic deinnervation seen in neurodegenerative disorders.

**Supplementary materials are attached after the main text.**

## Introduction

The cholinergic system arising from the basal forebrain (*1*, *2*) plays a fundamental role in controlling cortical functions including attention (*3*), learning and memory (*4*), plasticity (*5*), sleep–wake alternation (*6*), and is implicated in neurodegenerative diseases (*7*).

Contemporary models of the basal forebrain cholinergic system and efforts to develop pro-cholinergic treatments have been based largely on the assumption that cholinergic cells release only a single transmitter and it is released non-synaptically (*8*–*13*). The seemingly rare synapses on cholinergic fibers (see Supplemental Information) supported the concept of non-synaptic transmission. However, highly precise cholinergic transmission during reward and punishment (*14*), recordings of phasic release (*10*, *15*, *16*), and the dependence of hippocampal synaptic plasticity on the millisecond-scale timing of the cholinergic input (*17*) challenge this textbook model of non-synaptic transmission by cholinergic fibers.

Therefore, we hypothesized that all cholinergic terminals establish synapses. After immunolabeling, we analyzed the real incidence of synapses, localized vesicle pools using STORM super-resolution imaging and we also localized membrane-docked neurotransmitter vesicles using electron tomography. Because previous data suggested the co-localization of acetylcholine and GABA in retina and other brain areas (*18*–*23*), we also hypothesized that hippocampal cholinergic synapses may be GABAergic as well. Using immunolabeling and optogenetics combined with in vitro electrophysiology, we investigated the possible presence and subcellular regulation of hippocampal co-transmission of acetylcholine and GABA, and the role of its GABAergic component in controlling hippocampal network activity.

Challenging a decades-old model, we show that all hippocampal cholinergic terminals established synapses and vesicles dock only at synapses. We also show that hippocampal cholinergic synapses are GABAergic as well and evoke fast composite (hyperpolarizing and depolarizing) postsynaptic potentials. The two transmitters showed different short-term plasticity, were modulated by distinct calcium channels, and were mutually regulated by presynaptic auto-receptors. We demonstrate here that synaptic release of GABA from cholinergic terminals alone can suppress hippocampal sharp wave-ripples effectively and it can attenuate hippocampal epileptiform activity as well.

Our data urge the reinterpretation of previous studies about the basal forebrain cholinergic system and offer a new explanation for the emergence of hippocampal epileptiform activity associated with Alzheimer’s disease related loss of cholinergic innervation.

## Results

### All hippocampal cholinergic axon terminals form synapses, with GABA as a co-transmitter

Previous studies concluded that few cholinergic terminals form synapses (see Discussion and Supplemental Information). We hypothesized that all cholinergic terminals form synapses. By identifying cholinergic synapses with neuroligin 2 (NL2) labeling (Figure 1, S1, *24*), we could test the incidence of cholinergic synapses with three-dimensional serial electron microscopic reconstructions in the hippocampus. We identified septo-hippocampal cholinergic fibers with choline acetyltransferase (ChAT) labeling or using Cre-dependent eYFP-adeno-associated viruses (AAV) injected into ChAT-Cre mice (Figure 1, S1, S2; Supplemental Experimental Procedures). Having verified that all hippocampal cholinergic terminals originate from basal forebrain cholinergic cells (for controls see Supplemental Information and Figure S2), we found that all hippocampal cholinergic terminals examined established one or more NL2 positive synapses (Figure 1, S1; Table S4, Supplemental Information).

**Figure 1.**
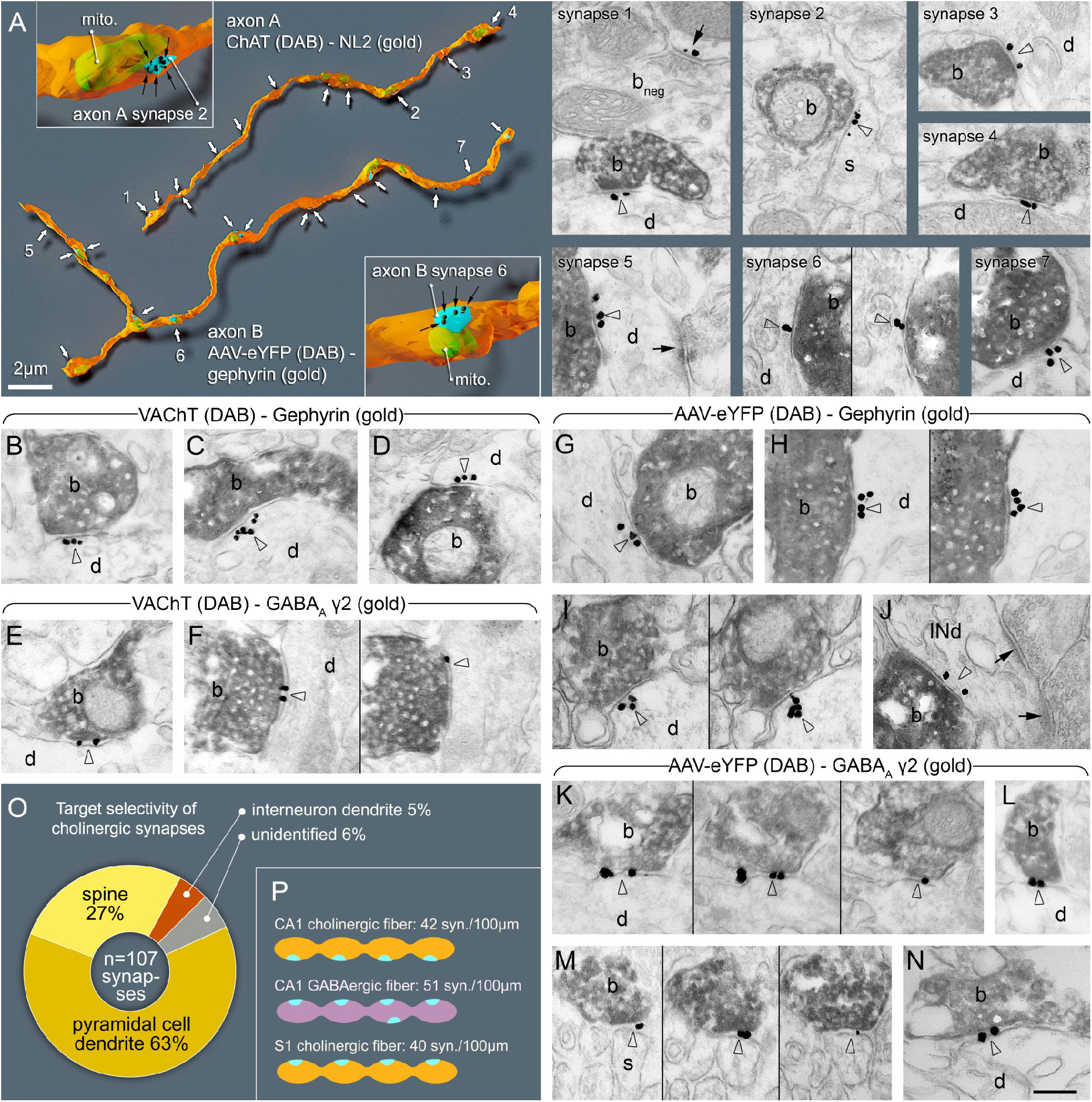
All cholinergic axon terminals establish synapses, and also express GABAergic markers postsynaptically, while innervating pyramidal cells and interneurons. **A:** Three-dimensional EM reconstructions show that hippocampal cholinergic fibers form synapses (arrows) frequently. Axon A is labeled for ChAT in a WT mouse. Axon B is an AAV-eYFP virus-labeled septo-hippocampal fiber in a ChAT-Cre mouse. Insets show two typical terminals with synapses (blue). The plasma membrane was made partially transparent to reveal mitochondria (mito, green). Gold labelings of NL2 (axon A, synapse 1-4) or gephyrin (axon B, synapse 5-7) were used to recognize synapses (black dots and arrows in the insets). EM images show terminal boutons (b) of the reconstructed axonal segments establishing synapses 1-7 (arrowheads, indicated by the same numbers on the left) on dendrites (d) and a spine (s). Next to synapse 1, a ChAT-negative, putative GABAergic terminal bouton (b_neg_) forming a NL2-positive synapse (arrow) is also shown. **B-N:** EM images reveal the presence of gephyrin (arrowheads, gold; B-D; G-J) and GABA_A_ γ2 receptor subunits (arrowheads, gold; E-F; K-N) postsynaptically in synapses established by vAChT-positive terminals in WT mice (B-F; DAB, b) or by AAV-eYFP-labeled septo-hippocampal terminals in ChAT-Cre mice (G-N; DAB, b). Images of consecutive sections are separated by thin black lines. Terminals innervate dendrites (d) or spines (s). In J, the postsynaptic target is an interneuron dendrite (INd) that receives type I synapses as well (arrows). Synapses are from str. ori. (A, B-E, G-J, L, M), str. rad. (K, N) and str. l-m (F). Scale bar is 200 nm for all EM images. **O:** Postsynaptic target selectivity of reconstructed cholinergic axonal segments from str. oriens and radiatum. **P:** Comparison of the number of synapses per 100 μm cholinergic axonal segments in CA1 and S1 cortex and GABAergic fibers in CA1.

Cholinergic synapses innervated pyramidal dendritic shafts (63%), spine-necks (27%), and interneuron dendrites (5%). Data from the somatosensory cortex was similar (Figure 1 O, Table S4; Supplemental Information). The linear density of synapses along cholinergic axons was similar to that found along GABAergic axons that are known to establish synapses abundantly (Figure S1, Table S4, and Supplemental Information).

Next, we tested whether these synapses are GABAergic as well. First, we localized the elements of postsynaptic GABAergic signaling machinery in these cholinergic contacts. We localized gephyrin first because it is known to interact with both GABA_A_ receptors and NL2. We found, that more than 81% of hippocampal cholinergic synapses contained gephyrin postsynaptically, on dendrites and spine-necks (Figure 1, Supplemental Information, Table S3). In addition, we found that more than 80% of the cholinergic synapses showed GABA_A_ receptor gamma2 subunit labeling that was also readily detected on both dendrites and spine-necks (Figure 1, S1, Table S3, and Supplemental Information).

Then, we localized the elements of presynaptic GABAergic and cholinergic signaling machinery in these terminals. By crossing a zsGreen fluorescent reporter-mouse-line with a vesicular GABA transporter (vGAT)-Cre mouse line and labeling the medial septum for ChAT, we found that all septo-hippocampal cholinergic cells are also vGAT positive (Figure 2A, B, Supplemental Information). Hippocampal cholinergic terminals expressed the GABA-synthesizing enzyme, glutamate decarboxylase 65 (GAD65) as well (Supplemental Information, Figure 2C). In addition, at least 83% of cholinergic septo-hippocampal terminals were vGAT positive (Figure 2D, Supplemental Information), whereas vesicular acetylcholine transporter (vAChT) was detected in 64% of the septo-hippocampal cholinergic terminals (Supplemental Information). Finally, using postembedding immunogold staining, we showed that GABA is detectable in cholinergic terminals (Figure 2 E-F, Supplemental Information).

**Figure 2.**
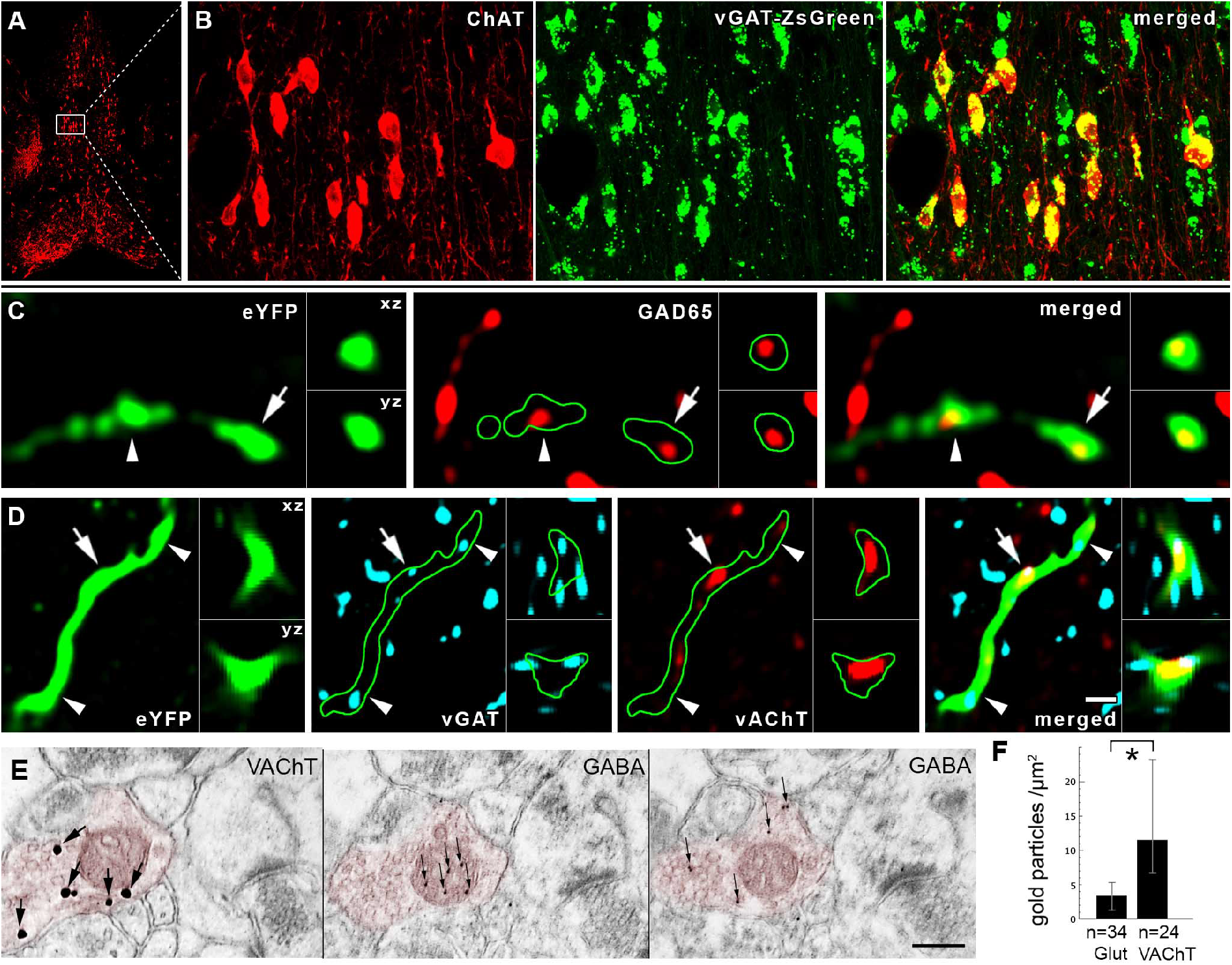
Cholinergic cells express the molecular machinery required for GABA release. **A-B:** The cholinergic neurons of the MS are GABAergic. White box in A contains area enlarged in B. Images show neurons stained for ChAT in red, while the green labeling marks the vGAT-expressing neurons in vGAT-ZsGreen reporter mouse. **C-D:** AAV-eYFP virus-traced septo-hippocampal fibers express GAD65 (C). AAV-eYFP virus-traced septo-hippocampal fibers express vGAT and vAChT (D). Insets show xz and yz projections of the terminal labeled with an arrow. Arrowhead points to another terminal. Green line marks the fiber outline. (Scale bar on D is 210, 14, 2 and 1 um for A, B, C and D, respectively.) **E-F:** Hippocampal cholinergic terminals contain GABA. Three consecutive EM sections of a vAChT-positive terminal (E, red pseudocolor) are shown. vAChT was visualized by preembedding immunogold method (the first panel of E, silver-intensified gold particles, large arrows), whereas on the next ultrathin sections (the second and third panels of E) postembedding GABA immunostaining was performed (smaller gold particles, thin arrows, some GABA molecules penetrate into mitochondria during fixation). vAChT signal is absent in postembedding images, because of the etching procedure. Scale bar is 200 nm for all EM images. **F:** Cholinergic terminals contained significantly higher immunogold signal (p< 0.05) suggesting the presence of GABA in terminals.

### Selective optogenetic activation of cholinergic axons evokes fast, composite GABAergic-cholinergic postsynaptic responses in the hippocampus

We investigated the electrophysiological properties of these cholinergic synaptic responses in target neurons. We injected Cre-dependent, channelrhodopsin 2 (ChR2) expressing adeno-associated virus (AAV-ChR2) into the medial septum of ChAT-Cre knock-in mice (Figure 3A, Supplemental Experimental Procedures) and recorded hippocampal cells. AAV-ChR2 labeled only cholinergic cells in the septo-hippocampal pathway (see Supplemental Information) and cells of wild type mice did not respond to illumination, proving that membrane potential responses in ChAT-Cre mice were caused only by cholinergic fibers. NMDA- and AMPA-type glutamate receptors were blocked to prevent polysynaptic recruitment of neuronal activity in all in vitro recordings presented in Figure 3 and 5.

**Figure 3.**
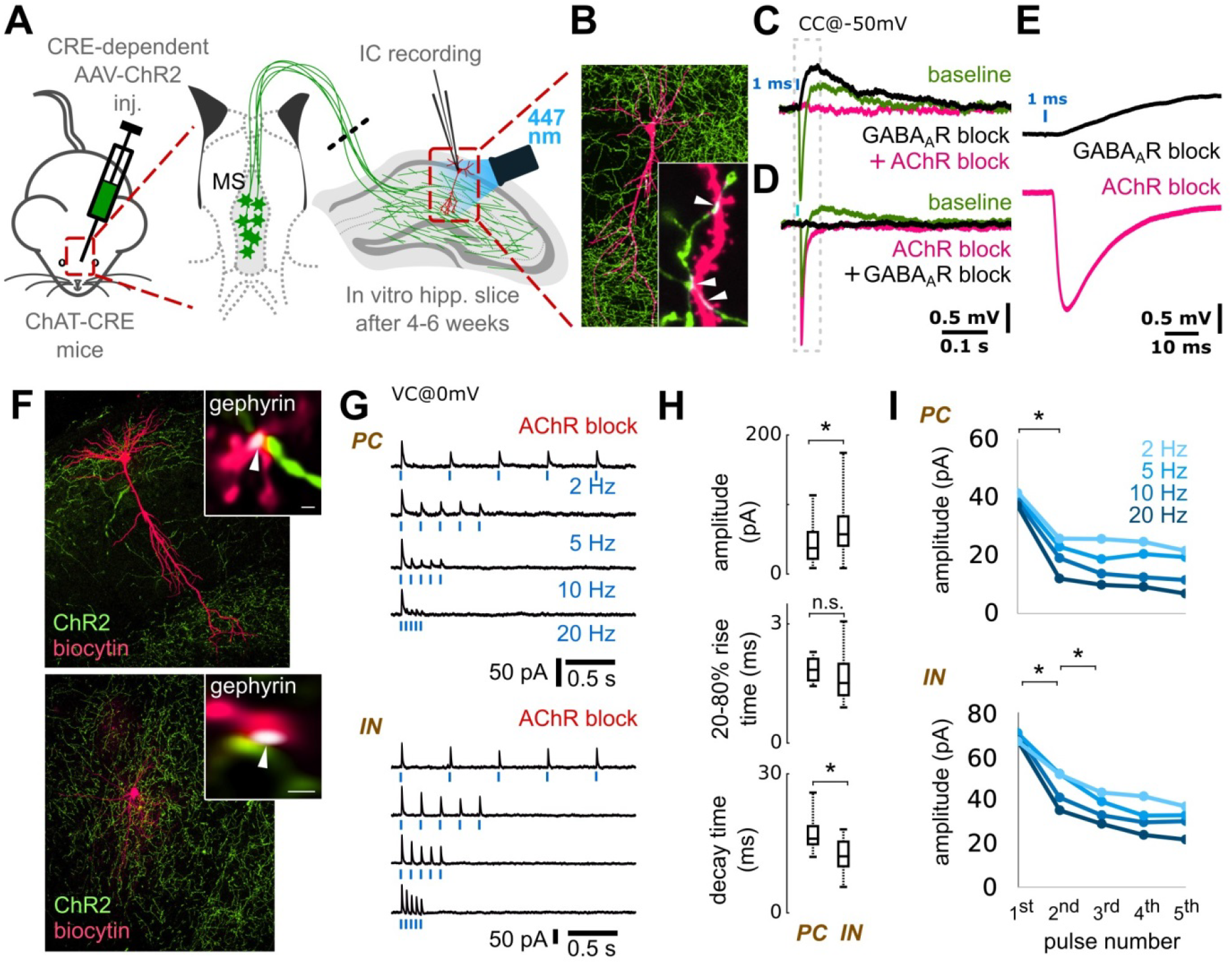
Optogenetic stimulation of cholinergic fibers elicits fast composite GABAergic and cholinergic postsynaptic responses. **A:** Medial septum (MS) of ChAT-Cre mice were injected with Cre-dependent AAV containing Channelrhodopsin-2 (ChR2). Using whole-cell patch clamp in horizontal hippocampal slices, we recorded voltage or current response from hippocampal neurons upon optical excitation of septo-hippocampal cholinergic fibers. NMDA and AMPA receptors were blocked with AP5 (50 μM) and NBQX (20 μM) in all experiments presented in Figure 3 and 5. **B:** Representative post-hoc visualized CA1 pyramidal cell (magenta) and the surrounding cholinergic fibers (green) with putative contacts (inset, white arrowheads). **C:** Blue light pulses elicited a composite membrane potential response from str. lacunosum-moleculare inhibitory neurons (green, average of 50 stimulations). Inhibition of GABA_A_Rs (10 μM gabazine) abolishes the hyperpolarizing component (9 from 9 tested cells), resulting in a putative cholinergic excitatory potential (black). Inhibition of all AChRs (1 μM MLA, 1 μM DHβE, 10 μM atropine, 4 from 4 tested cells) blocks the remaining depolarizing response (magenta). **D:** Conversely, first blocking the AChRs, resulted in a putative GABAergic IPSP (magenta), which was blocked by GABA_A_Rs inhibitor gabazine (black). The increase of IPSP amplitude for AChR block is addressed in Figure 5. **E:** Magnified cholinergic EPSP (black) from panel C, and GABAergic IPSP from panel D (magenta) demonstrate their short latency (see also Figure S3). **F:** A representative recorded pyramidal cell (PC, top) and an inhibitory neuron (IN, bottom) post-hoc visualized in magenta, and ChR2 positive cholinergic axons in green. **Insets:** Immunostaining for gephyrin (white) identify their putative synapses (white arrows). **G:** We blocked both nicotinic and muscarinic AChRs (1 μM MLA, 1 μM DHβE, 10 μM atropine) and recorded inhibitory postsynaptic currents (IPSC) evoked by cholinergic fiber illumination. Traces show IPSC response of a PC and an IN to 5 light pulses (1ms) at increasing stimulation frequencies (2, 5, 10, 20 Hz, cells were recorded in VC@0mV). **H:** Amplitude and decay time of unitary GABAergic IPSCs from pyramidal cells (n=5) and inhibitory neurons (n=16) were statistically different (p<0.05), while their rise time was not different (see Supplemental Information) **I:** Averages of IPSC amplitudes for the 5 pulses presented on panel G shows strong short-term depression (STD) of GABAergic transmission evoked by stimulating cholinergic fibers (for details see Supplemental Information and Figure S3).

**Figure 4.**
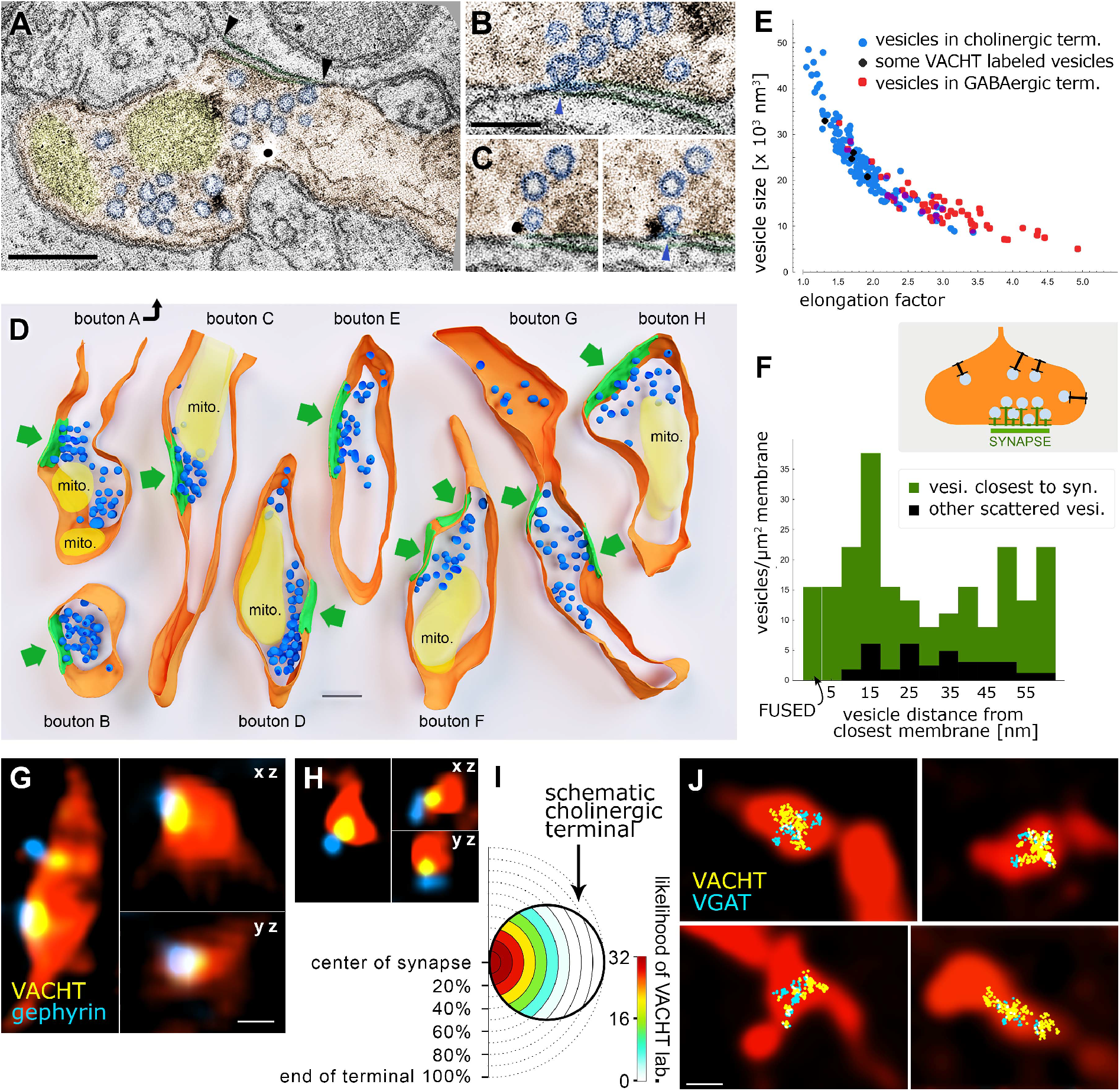
Separate vesicular release of acetylcholine and GABA originate from the same synaptic active zones in cholinergic fibers. **A-C:** About 0.5 nm thick virtual sections from electron tomographic reconstructions of cholinergic terminals. Terminal is orange, mitochondria are yellow, synaptic vesicles are blue. Black arrowheads point to the edges of the synapse. Scale bar is 200 nm. **B:** Active zone is enlarged from A. The omega-shaped vesicle is being fused with the terminal membrane (blue arrowhead). **C:** Two serial virtual sections showing a portion of the active zone of another terminal. A fused synaptic vesicle (blue arrowhead) is labeled for vesicular acetylcholine transporter (black immunogold particle). Scale bar for B and C is 100 nm. **D:** Three-dimensional models of reconstructed terminals. Terminal membranes are orange, mitochondria are yellow, synaptic vesicles are blue and synapses are green. Green arrows point to the synapses. Scale bar is 200 nm. The association of the vesicle pools to the active zones is evident. **E:** Scatterplot shows the relationship of vesicle size (volume) and elongation factor in the sampled vesicles. Vesicles in the cholinergic terminals had a broader size-distribution than vesicles of purely GABAergic terminals, which were smaller and more elongated (Supplemental Information). Some vesicles (black markers) could be identified as vAChT-positive because immunogold particles clearly labelled them. **F:** The density of the vesicles within a 60-nm-wide band from the membrane is plotted as a function of distance from the terminal membrane at synaptic (green) and extrasynaptic (black) areas. Vesicle density was 6.5 times higher (in the first 60 nm from the membrane) in the synaptic active zone than extrasynaptically (n=183 vesicles from 8 terminals in 2 mice). Docked (closer to the membrane than 5 nm) or fused (undergoing exocytotic fusion) vesicles were absent extrasynaptically, but were present in the active zones. **G-H:** vAChT labeling was confined to synaptic active zones in cholinergic fibers. CLSM images show terminals of virus-traced septo-hippocampal fibers (red pseudocolor) containing vAChT-positive vesicle pools (yellow), localized to the synaptic active zones that are identified by gephyrin labeling (blue). Scale bar is 500 nm for G and H. **I:** Scale-free analysis confirms that the likelihood of vAChT labeling is the highest at the synaptic active zones (n=32 synapses, 2 mice). **J:** Correlated CLSM-STORM super-resolution microscopy images show that vAChT (yellow) and vGAT (cyan) vesicle pools are overlapping and are confined to small portions within AAV-eYFP virus-traced septo-hippocampal fibers (red pseudocolor).

**Figure 5.**
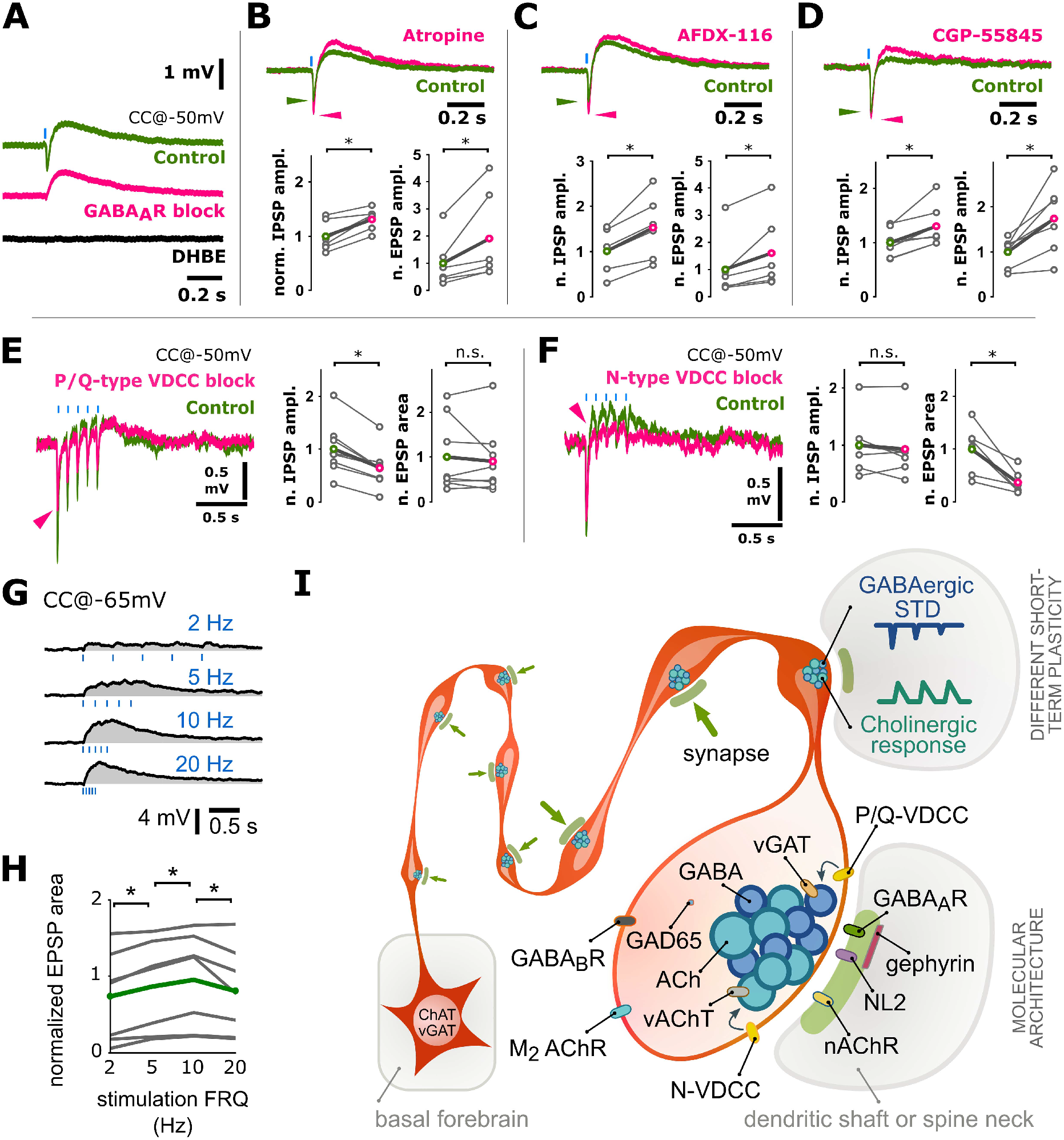
Co-transmission of GABA and acetylcholine is mutually cross-regulated by autoreceptors, mediated by distinct calcium channels and shows different short-term plasticity. **A:** Optical stimulations of cholinergic fibers trigger composite membrane potential responses (average of 50 traces) of an inhibitory neuron recorded in str. lacunosum-moleculare (green). Cells were held at approximately -50 mV to record hyperpolarizing and depolarizing components concurrently. Early hyperpolarization was confirmed to be of GABAergic origin by bath application of the GABA_A_R blocker gabazine (magenta, 10 μM, which abolished quick hyperpolarization in 9 out of 9 tested cells), while the late depolarization was confirmed to be cholinergic by the application of the alpha4-type nicotinic acetylcholine receptor blocker DHβE (black, 1 μM, which abolished depolarization in 5 out of 5 tested cells). Cells with identifiable nicotinic responses were used to further investigate the properties of GABA and acetylcholine release from cholinergic fibers. **B:** Increase in GABAergic response after AChR block (Figure 3D) suggested that presynaptic muscarinic receptors cross-modulated GABA (and possibly ACh) release from cholinergic fibers. Therefore, putative presynaptic muscarinic receptors were selectively inhibited via bath application of atropine (n=6), which led to a significant increase in amplitude of both the GABAergic hyperpolarization and nicotinic depolarization (green, see Supplemental Information). **C:** Bath application of the selective M2 antagonist AFDX-116 (10 μM, n=6) resulted in changes similar to that of atropine application (for data see Supplemental Information). **D:** Blocking GABA_B_Rs with CGP-55845 (1 μM, n=7) also resulted in increased GABAergic and cholinergic responses suggesting a dual modulatory role (for data see Supplemental Information). **E-F:** Testing roles of distinct voltage-dependent calcium channels (VDCC) in release of GABA and Ach **E**: Blockade of P/Q-type VDCC, using selective antagonist ω-agatoxin (1 μM, puff application), reliably decreased GABAergic response component but did not alter cholinergic component. At control conditions, 5 short (1 ms) light pulses at 10 Hz evoked a composite event of GABAergic hyperpolarization and cholinergic depolarization (green, average of 20 traces). In response to ω-agatoxin application (magenta, n=9), the GABAergic component (measured as IPSP amplitude) decreased robustly (magenta arrowhead), and the cholinergic component (measured as EPSP area) showed no change (for details see Supplemental Information). **F:** Blockade of N-type VDCC, using selective blocker ω-conotoxin (1 μM), reliably reduced cholinergic response component (n=6) but left GABAergic component unchanged (for details see Supplemental Information). **G:** Short-term dynamics of cholinergic EPSPs recorded from inhibitory neurons in str. lac.-mol. in response to 5 light pulses at different frequencies (top; 2, 5, 10, 20 Hz, n = 7). Cholinergic events overlap, not allowing reliable EPSP peak detection. Therefore, STP was quantified as the integral of the evoked events. **H:** The relative change in EPSP integral for the stimulations at different frequencies (n=7, average is green). Unlike the GABAergic component (Figure 3), short-term depression was not observed in cholinergic responses. Slight increase in EPSP area from 2 to 10 Hz suggests some form of a weak facilitation of the cholinergic component. **I:** Summary illustration of some of the findings. All cholinergic terminals establish synapses, they are fully equipped with GABAergic-cholinergic co-transmission signaling machinery. GABAergic and cholinergic vesicles are regulated by different VDCCs. GABAergic transmission shows strong STD, while the cholinergic one is rather facilitating or not changing. Synapses are established on both dendritic shafts and spines in hippocampus.

We recorded the membrane potential of inhibitory neurons in CA1 str. lacunosum-moleculare, because they are known to display cholinergic responses (*25*). Cells were recorded using whole-cell patch-clamp in responses to 1ms optical stimulation (Figure 3A, Supplemental Experimental Procedures) that resulted in a composite membrane potential response: a quick GABA_A_ receptor-dependent hyperpolarization (peak @ 13.8 ms), and a slightly delayed (peak @ 92 ms) acetylcholine receptor-dependent depolarization (Figure 3C-E, S3A-C). Although these synaptically released transmitters may act on non-synaptic receptors as well, both responses had short onset latency (Figure S3A-C), which together with our anatomical data suggests synaptic release and action of GABA. Synaptic spill-over of GABA and acetylcholine may act extrasynaptically as well.

Next, we blocked both nicotinic and muscarinic AChRs and recorded inhibitory postsynaptic currents (IPSC) on pyramidal cells (PCs) and interneurons (INs) after optical stimulation of cholinergic fibers (Figure 3F-I). Single IPSC kinetics and short-term plasticity in PCs and INs were tested using five short light pulses at physiologically relevant firing rates (at 2, 5, 10 and 20 Hz) measured in vivo (*26*, *27*). The amplitude of the evoked inhibitory currents (calculated for the first stimulus) was larger on INs than on PCs, but their rise-time (20-80%) was not significantly different. IPSCs evoked in INs had a shorter decay time (Figure 3H). The series of light pulses revealed strong short-term depression (STD) of inhibitory currents onto both PCs and INs (Figure 3G and I). GABAergic STD was observed in every tested neuron and was accompanied by a decrease in transmission probability during the stimulation sequence (Figure S3), suggesting a presynaptic mechanism for STD.

Calcium influx through ChR2 expressed on axon terminals can alter intrinsic short-term plasticity of the synapses (*28*). To exclude this possibility, we used a digital micromirror device (DMD) inserted into the optical path of the microscope, to illuminate cholinergic axons running towards the recorded cell, but not the terminals (Figure S3D). This resulted in STD similar to that recorded with optic fiber illumination (Figure S3E-F), excluding the possibility of ChR2 mediated Ca^2+^ influx, as the reason for the observed STD. A series of stimuli could also decrease the driving force of chloride through GABA_A_Rs, resulting in apparent STD (*29*, *30*). In this case, the putative site of STD would be postsynaptic instead of presynaptic. When we used a high Cl^-^ intracellular solution to prevent shifts in Cl^-^ reversal, a series of stimuli resulted in STD similar to that shown above (Figure S3G and H).

### Cholinergic and GABAergic vesicle docking is restricted to synaptic active zones in cholinergic fibers

Our physiological and anatomical data showed that cholinergic terminals establish effective cholinergic-GABAergic synapses. Non-synaptic acetylcholine release, however, might still be possible if cholinergic vesicles could be docked and fused outside of the synaptic active zones as well. Using advanced electron tomography tools, we were able to reconstruct cholinergic fiber segments, to localize vesicles with 1 nm resolution and to analyze the precise distribution of synaptic vesicles in hippocampal cholinergic terminals. We identified terminals using vAChT immunogold labeling and reconstructed them using electron tomography (Figure 4A-D). Three-dimensional reconstruction revealed that synaptic vesicles clustered close to the active zones (Figure 4D). We measured the distances of vesicles to the closest (synaptic or non-synaptic) cell membranes, and compared their density at different distance intervals from the membranes (Figure 4F). The density of the vesicles within 60 nm from the membranes was 6.5 times higher in the synaptic active zone than extrasynaptically (Figure 4D, F). We did not find any docked (less than 5 nm from the membrane) or fused (undergoing exocytotic fusion) vesicles non-synaptically, but we detected both of these at synapses (Figure 4B, C, F). The distribution of vesicles in cholinergic terminals was similar to that found at other glutamatergic or GABAergic terminals, arguing against non-synaptic vesicular docking or release in cholinergic terminals.

Vesicular volume can correlate with its transmitter content (*31*). Therefore, using electron tomography, we compared the vesicular morphology between cholinergic and GABAergic terminals (Figure 4E). Vesicles of cholinergic terminals were significantly (p<0.001), about 60% larger than those in GABAergic terminals (Figure 4E, Supplemental Information) and their volumes were significantly more variable as well (p<0.001). Smaller (putatively GABAergic) vesicles of cholinergic terminals were similar to those in purely GABAergic terminals (Figure 4E, see Supplemental Information), suggesting an even larger difference between the two types of vesicles.

### Acetylcholine and GABA are released at the same synapses, but from different vesicles in cholinergic terminals

To further confirm that the two transmitter systems use the same active zones, we used confocal fluorescent imaging in mouse hippocampus and found that vAChT-positive vesicles were concentrated at gephyrin-labeled, putative GABAergic-cholinergic synapses (Figure 4G, H, Supplemental Information). Scale-free analysis confirmed that the likelihood of vAChT labeling was the highest at the gephyrin-labeled synaptic active zones (Figure 4I, Supplemental Information).

Using super-resolution STORM imaging of vGAT and vAChT immunolabeling, combined with correlated fluorescent confocal laser scanning microscopy (CLSM) of cholinergic fibers, we demonstrated that cholinergic-GABAergic vesicle pools were mixed and were confined to a small volume of the AAV-eYFP labeled septo-hippocampal terminals (Figure 4J, for details, see Supplemental Information).

Using isolated mouse cortical synaptic vesicles, we found that acetylcholine and GABA are packed into different vesicles (Figure S4A, Supplemental Information, and Supplemental Experimental Procedures). Both flow cytometry (Figure S4B) and electron microscopy data (Figure S4C, E) confirmed the purity of our sample (see also Supplemental Information and Experimental Procedures). All vesicles were labeled with synaptophysin (SYP). After quadruple-immunolabeling of the isolated synaptic vesicles, 29% were only SYP-positive, 45% were double-labeled for vesicular glutamate transporter 1 and SYP, 14% were double-labeled for vGAT and SYP, 11% were double-labeled for vAChT and SYP. Only a negligible amount of vesicles (0.9%) were triple labeled with any combinations and only 0.14% of all vesicles were co-labeled for cAChT and vGAT, suggesting that vesicular transporters for GABA and acetylcholine are expressed by distinct vesicle populations in cortical samples (Figure S4H; Supplemental Information).

### Acetylcholine and GABA release shows different short-term plasticity, modulated by distinct voltage-dependent calcium channels, and are mutually regulated by different presynaptic auto-receptors

Our anatomical data demonstrated that acetylcholine and GABA are released from different vesicles. However, distinct vesicle pools of the same terminals can couple to different release mechanisms with different short-term plasticity (*20*). Because GABAergic transmission always show strong STD in cholinergic terminals, we examined, whether cholinergic responses display similar short-term dynamics. We recorded inhibitory neurons and light-stimulated ChR2 expressing cholinergic axons (Figure 5G). Their net depolarization for a series of five stimuli did not decrease but increased slightly with frequency (Figure 5H). We found that cholinergic dynamics did not show short-term depression present in GABAergic responses, in which case net depolarization should have decreased. The observed slight increase in the net depolarization with increasing stimulation frequency indicates a form of facilitation in cholinergic responses. This differential dynamics of GABA and acetylcholine release confirmed that the two transmitters are released from distinct vesicular pools.

Dynamics of vesicle release likely depends on the type of voltage-dependent calcium channels (VDCCs), which mediate Ca^2+^ triggered vesicle release (*32*). However, different VDCCs in the same cell (*20*) can be associated with distinct types of vesicles. Indeed, we found that selective blockade of P/Q-type calcium channels decreased GABAergic IPSPs significantly, but caused no change in the cholinergic components (Figure 5E, Supplemental Information). Conversely, the cholinergic component was robustly decreased after the selective blockade of N-type calcium channels, while GABAergic IPSPs showed no change (Figure 5F, Supplemental Information). Besides confirming the presence of different vesicular pools, this revealed different molecular pathways for their regulation during co-transmission.

We also investigated presynaptic modulation of this vesicular release. Blocking AChRs increased the amplitude of GABAergic hyperpolarization (Figure 3D), for which presynaptic muscarinic receptors may be responsible (*25*). We held CA1 interneurons at -50 mV, to record GABAergic hyperpolarization and cholinergic depolarization concurrently (Figure 5A). Muscarinic receptor blocker atropine (10 μM) significantly increased the amplitude of both GABAergic IPSPs and cholinergic EPSPs (Figure 5B). By blocking M2-type AChRs, reported abundant in hippocampal cholinergic terminals (*25*, *33*), we reproduced PSP increases evoked by atropine (Figure 5C). This confirms the role of M2-type AChRs in regulating both acetylcholine and GABA release from cholinergic terminals.

Previously, we described the presence of presynaptic GABA_B_ autoreceptors in septal cholinergic cells (*34*). To test their role in regulating synaptic co-transmission, we blocked GABA_B_ receptors. This led to a significant increase in the amplitude of both GABAergic IPSPs and cholinergic EPSPs (Figure 5D). However, we did not see GABA_B_ receptor-dependent postsynaptic hyperpolarization in response to cholinergic fiber stimulation in our recordings. Overall, these experiments revealed that GABA and acetylcholine cross- and auto-regulate their co-transmission from cholinergic-GABAergic terminals, presynaptically.

### GABA release from septal cholinergic cells is essential for shaping healthy and pathological cortical network activities

Septal cholinergic neurons control hippocampal activity states (*35*, *36*) and suppress *in vivo* sharp wave-ripples (SWR) in the hippocampus (*37*). However, the effect of their hippocampal GABA release has not been investigated. We recorded spontaneous SWRs in hippocampal slices (Figure 6, Supplemental Experimental Procedures). Illumination of control slices without ChR2 expression caused no change in SWR activity. In AAV-ChR2 infected ChAT-Cre mice, we found that without blocking cholinergic transmission, optical stimulation of cholinergic fibers decreased SWR rate significantly (Figure 6C-D), followed by a transient increase in SWR rate after the cessation of optical stimuli. However, when cholinergic transmission was blocked, the same optical stimulation caused a similar decrease in SWR rate (Figure 6E-F), with no rebound after stimulation. Although cholinergic fibers have been reported to be responsible for inhibiting SWRs (*37*), here we demonstrated that GABA release from cholinergic synapses alone is sufficient to downregulate the rate of SWRs.

**Figure 6.**
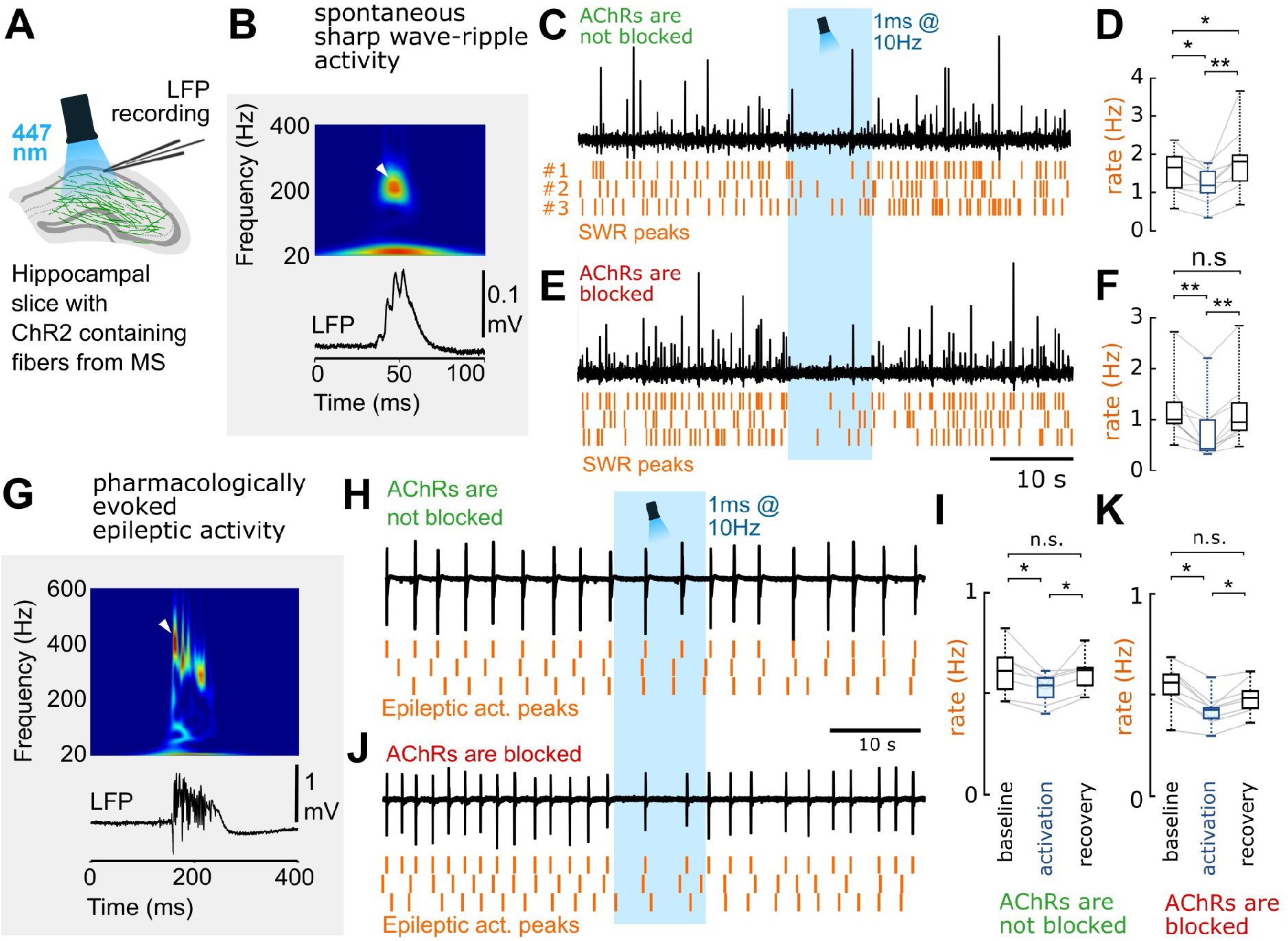
GABA release from cholinergic terminals effectively controls healthy and pathological hippocampal network activity. **A:** To understand the functional impact of GABA and acetylcholine co-transmission onto cortical network oscillations, local field potential (LFP) was recorded with standard patch pipettes from the CA1 area of thick hippocampal slices. These slices generated healthy and pathological in vivo like activity patterns. Cholinergic fibers expressing ChR2 were illuminated with blue light in the presence, or in the absence of AChR blockers parallel with LFP recordings. **B:** A spontaneous sharp wave-ripple (SWR) LFP signature (bottom), and its wavelet transform (top) highlighting the characteristic ripple-band frequency component (~200 Hz, white arrowhead). **C-D:** LFP recording from hippocampal CA1 in the absence of AChR blockers (n=9). Orange raster indicate SWR peaks for three consecutive stimulation periods. Optical stimulation decreased SWR rate significantly (D), then SWR rate increased temporarily after the cessation of optical stimulation (for data see Supplemental Information). **E-F:** After we blocked AChRs (1 μM MLA, 1 μM DHβE, 10 μM atropine), LFP recording from CA1 (n=9) showed that GABA release from cholinergic fibers alone could decrease SWR rate even more effectively (F). There was no SWR rate increase after the cessation of optical stimulation (for data see Supplemental Information). **G:** Epileptiform activity was evoked by elevating K^+^ concentration in the ACSF to 8.5mM in slices. LFP (bottom) and its wavelet transform (top) shows the signature of a pharmacologically evoked epileptiform events. Note the robust differences in amplitude, length and frequency components compared to physiological SWR activity (Figure 6B) that is consistent with literature data (*39*). **H-I:** Epileptic activity recorded in CA1 in the absence of AChR blockers (n=7). Orange raster indicate detected epileptic activity peaks for three consecutive stimulation periods. Illumination of cholinergic fibers reduced the rate of epileptic events (I), which recovers after the cessation of optical stimulation (for data see Supplemental Information). **J-K:** Epileptic activity recorded in CA1 in the presence of AChR blockers (n=8). Even in the presence of AChR blockers, illumination of cholinergic fibers reduced the rate of epileptic discharges (K), which recovers after the stimulation (for data see Supplemental Information), suggesting a crucial role for GABA release in controlling epileptiform activity.

The GABA release from septo-hippocampal cholinergic fibers may also affect the occurrence of hippocampal epileptiform activity. Therefore, we examined the effect of GABA release from cholinergic axons in the in vitro high potassium epilepsy model (Figure 6G-K, *38*, *39*). Optical stimulation of cholinergic fibers reduced the rate of epileptiform events (Figure 6H-I) and a similar reduction was present, when AChRs were blocked (Figure 6J-K). These results demonstrated that synaptic co-transmission of GABA from cholinergic fibers alone can effectively control hippocampal epileptiform activity.

## Discussion

Our results challenge the model of non-synaptic single transmitter release from cholinergic fibers in the hippocampus and demonstrate a synaptic communication using two neurotransmitters. Our results also show non-synaptic action of these synaptically released transmitters. We showed that hippocampal cholinergic synapses were effective GABAergic synapses as well. Cholinergic fibers are among the densest and most influential subcortical pathways in hippocampus; therefore, our findings suggest a fundamental change in our view of the regulation of hippocampal states. We also found that acetylcholine and GABA were not co-released but co-transmitted from the same synapse. We confirmed this by showing that (i) isolated cortical transmitter vesicles do not co-express vGAT and vAChT, (ii) GABAergic and cholinergic components show different short-term plasticity in these fibers, (iii) they are preferentially modulated by distinct voltage-dependent calcium channels, and (iv) electron tomography suggested differences in the volume of GABA and acetylcholine containing vesicles. This co-transmission (similar to that in retina *20*) may require nanoscale subsynaptic organization of presynaptic molecules, as proposed before (*40*). Emphasizing the functional role of GABA release, we also demonstrated that synaptic GABA release from cholinergic terminals alone can effectively suppress hippocampal sharp wave-ripples and *per se* can attenuate hippocampal epileptiform activity. These confirmed the functional importance of this GABAergic-cholinergic co-transmission in healthy and pathological states and led to a novel model of the septo-hippocampal cholinergic neurotransmission.

### Cholinergic terminals establish synapses

For decades, the predominant form of cholinergic communication was thought to be a form of “non-synaptic transmission" (*8*–*10*, *12*, *41*), which was seemingly supported by studies showing cholinergic fibers with few synapses (for details see Supplemental Information). Originally, the presence of extrasynaptic acetylcholine receptors and micro-dialysis experiments seemed to support a non-synaptic transmission hypothesis; however, later, sensitive microelectrodes showed faster changes in extracellular acetylcholine levels (Supplemental Information). In addition, basal forebrain cholinergic neurons were also shown to respond to reward and punishment with extremely high speed and precision (*14*), and recent data suggested that cholinergic cells may regulate cortical information processing with a remarkable, millisecond-scale temporal precision (*15*, *17*, *42*, *43*). These data urged the re-examination of whether acetylcholine release is synaptic (*44*). Using electron tomography and direct labeling of cholinergic synapses using neuroligin 2, we showed that, all terminals of the dense meshwork of hippocampal cholinergic fibers established (one or more) synapses and no docked or fused vesicles were detectable non-synaptically. Most of these synapses were missed previously, probably, because of their weak membrane thickening and narrower synaptic gap. Furthermore, we found that vAChT labeled vesicle pools associated only with synaptic active zones. Although synaptically released acetylcholine or GABA may act on non-synaptic receptors as well, in a “volume transmission” fashion, none of our data supports non-synaptic release. Therefore, the most likely resolution for the mismatch between this and the classic non-synaptic volume transmission hypothesis is a strong “spill-over” of transmitters from these frequent synapses that would allow transmitters to reach extrasynaptic receptors as well. In fact, the much larger cholinergic synaptic vesicles that we found could contain more acetylcholine to counteract its effective extrasynaptic removal by extracellular acetylcholine esterase (AChE).

### The tight regulation of synaptic GABAergic co-transmission from basal forebrain cholinergic fibers and their role in hippocampal synaptic plasticity

Co-transmission can significantly increase the efficacy of information transfer (*45*–*47*). Using purified vesicle preparation, we showed that GABA and acetylcholine are not “co-released” from the same vesicles, but “co-transmitted” from separate vesicles, sequentially inhibiting and exciting hippocampal neurons. To unlock the full potential of co-transmission, these coexisting vesicle pools need complex regulation. Indeed, we showed that the release of GABAergic vesicles are preferentially regulated by P/Q-type calcium channels, while the cholinergic ones are more affected by N-type calcium channels in the same synapses, similar to that of retinal co-transmission of GABA and acetylcholine (*20*). Differential calcium channel regulation may be achieved by sub-synaptic nanoscale organization of presynaptic molecules (*40*) providing partly different access for the distinct vesicle pools. Negative feedback via presynaptic autoreceptors could provide a further level of control. Previously, we demonstrated that septo-hippocampal cholinergic neurons express GABA_B_-receptors (*34*), while the expression of muscarinic M2 receptors was also described in these cells (*33*). Indeed, we confirmed that blocking either M2- or GABA_B_-receptors increased the release of both acetylcholine and GABA, suggesting the presence of tonically active autoreceptor regulation of these transmitters, presynaptically. In addition, our results, verified by a series of control experiments, suggested that GABAergic STD is an intrinsic property of synapses established by cholinergic fibers in the mouse hippocampus.

The timing of this co-transmission seems crucially important in hippocampal Schaffer collateral (SC) to CA1 synaptic plasticity. If the cholinergic input was activated 100 or 10 ms before SC stimulation, it resulted in ionotropic AChR-dependent long-term potentiation (LTP) or short-term depression, respectively, while delaying cholinergic stimulation until 10 ms after SC stimulation resulted in metabotropic AChR-dependent LTP (*17*). Here, we found that the GABAergic and cholinergic components of this co-transmission reach their IPSP and EPSP peak about 13.8 and 92 ms after stimulation, respectively, which suggest that synaptic plasticity may depend on the coincidence of SC stimulation with the GABAergic or cholinergic component of PSPs from these basal forebrain fibers. Nevertheless, the GABAergic component seems to have its own inhibitory role in the target network, making basal forebrain cholinergic fibers an unorthodox but effective source of GABAergic inhibitory control of the hippocampus, as suggested by our demonstration of its effect on network dynamics.

### Role of basal forebrain cholinergic system in regulating hippocampal network states

Basal forebrain cholinergic cells play a pivotal role in transforming activity states in the hippocampus (*35*). High cholinergic cell activity is associated with theta oscillation and the fast, yet unstable storage of external information in the hippocampus, while low cholinergic activity accompanies sharp wave-ripple (SWR) activity (*48*), which seems important for the consolidation and relocation of unstable memory traces from the hippocampus to the neocortex (*49*, *50*). Classic theories of cholinergic modulation presume that diffusely released acetylcholine would slowly retune cortical network activity, enabling the appearance of distinct network dynamics (*8*, *51*, *52*). Indeed, a recent study demonstrated that medial septal cholinergic cells suppress SWR activity in vivo (*37*). The authors suspected an M2 cholinergic receptor-mediated suppression of GABAergic interneurons. We re-examined that hypothesis and confirmed that cholinergic fiber activation indeed suppressed SWRs; however, we also showed that GABA (without acetylcholine) release from cholinergic fibers alone was able to achieve that. This GABAergic inhibition could lower the probability of concurrently active pyramidal cells in a given time-window, which is required for stochastic SWR initiation (*53*). Blocking all GABA receptors would have eliminated SWRs (*54*), therefore the effect of acetylcholine release alone, without the contribution of GABA co-transmission could not be addressed. The presence of a transient increase in SWR rate after the cessation of optical stimuli, might reflect longer-lasting changes in network or cell excitability, mediated by “synaptic spill-over” of acetylcholine, while the effect of GABA release from smaller vesicles is likely confined to synaptic (and presynaptic) receptors. However, this effect was not reported in vivo (*37*), therefore, it could have been the result of a relatively low AChE activity in our dual-superfusion system.

Degeneration of the cholinergic system is a typical characteristic of Alzheimer’s disease pathology (*7*), and these patients often develop epileptic seizures (*55*). Indeed, selective lesion of medial septal cholinergic projection increases seizure incidence in the hippocampus (*56*–*58*), while REM-sleep, which is associated with enhanced cholinergic cell activity (*27*), has a suppressing effect on epileptic seizures (*59*, *60*). The mechanisms, however, were unclear, because several cholinergic agonists were shown to trigger epileptic discharges (*61*). Therefore, we hypothesize that degeneration of basal forebrain cholinergic cells leads to epileptiform activity primarily, because it deprives the hippocampus of a massive GABAergic input. Indeed, our results supported this idea, because GABA release from basal forebrain cholinergic fibers alone was sufficient to decrease the occurrence of epileptiform activity in the hippocampus.

Our results, showing a tightly regulated, effective, synaptic GABA co-transmission from hippocampal cholinergic fibers urge the re-interpretation of previous models and can lead to alternative pharmacotherapies to treat Alzheimer’s disease-related loss of cholinergic innervation.

## Acknowledgements

This work was supported by the European Research Council (ERC-2011-ADG-294313, SERRACO), National Institutes of Health, USA (NS030549), National Research, Development and Innovation Office, Hungary (OTKA K119521, OTKA K115441 and VKSZ_14-1-2O15-0155), Human Brain Project, EU (EU H2020 720270).; B.P. is supported by UNKP-16-2-13 and D.S. is supported by the UNKP-16-3-IV New National Excellence Program of the Ministry of Human Capacities, Hungary. Á.D. is supported by the Hungarian Brain Research Program KTIA_13_NAP-A-I/2, the ‘Momentum’ Program of the Hungarian Academy of Sciences and ERC-CoG 724994. We thank Lászlé Barna, the Nikon Microscopy Center at IEM, Nikon Austria GmbH and Auro-Science Consulting Ltd for technical support for fluorescent and STORM imaging. We thank Emőke Szépné Simon, Katalin Lengyel, Katalin Iványi, Győző Goda and Nándor Kriczky for assistance and Anna Jász for help in scripting. We thank Dr. David Mastronarde (Univ. of Colorado Boulder) for help with electron tomography software IMOD. We thank Drs Lászlé Acsády, Julian Budd, Balázs Hangya, Liset Menendez de la Prida for their comment on an earlier version of the manuscript.

## Author Contributions

Conceptualization, V.T., C.C., D.S., G.N.; Methodology, V.T., C.C., D.S., G.N., Z.K., A.D, A.I.G.; Investigation, V.T., C.C., D.S., B.P., A.S., K.E.S., Z.K.; Writing – Original Draft, V.T., C.C., D.S., G.N.; Writing – Editing, V.T., C.C., D.S., G.N, A.D., A.I.G, T.F.F.; Funding Acquisition and Supervision, T.F.F., G.N., A.I.G., A.D.

## SUPPLEMENTARY MATERIALS FOR

### SUPPLEMENTARY RESULTS

#### All hippocampal cholinergic axon terminals form synapses that can be identified using NL2 labeling

We reconstructed randomly selected, long axonal segments [6 – 33 μm, average: 21 μm, n= 17, labeled either with anti-choline acetyltransferase (ChAT) antibody in wild type (WT) mice or with eYFP-adeno-associated viruses (AAV) injected in ChAT-Cre mice, Figure 1, S1, Table S4, for controls see Suppl. Experimental Procedures] and identified their synapses with NL2 or gephyrin immunogold labeling. All of them established synapses abundantly (Figure 1, S1). The average density of synapses was 42 synapses/100 μm. Some of these contact sites would not have been considered synapses earlier, because of their weak membrane thickening and narrower intercellular synaptic gap (e.g. Figure 1A, synapse 2-3) (*1*); however, NL2 and gephyrin labeling clearly identified their active zones. For comparison, we have also reconstructed GABAergic axonal segments (labeled for cannabinoid receptor type 1, CB_1_; n=2, 18 and 29 μm, Figure S1, Table S4), which are known to establish synapses abundantly. We found that practically all cholinergic terminals formed one or sometimes two synapses (Figure 1 and Figure S1). As a consequence, the linear density of synapses along cholinergic axons was similar to that along GABAergic axons (Figure S1, Table S4, number of synapses per 100 μm cholinergic axons: 42 in CA1, 40 in S1, number of synapses per 100 μm CB_1_-positive axons: 51).

#### Postsynaptic targets of cholinergic axons

Using electron microscopy in hippocampal CA1, we found that NL2 and gephyrin positive cholinergic synapses (n=107, collected from 4 mice) predominantly innervated pyramidal dendritic shafts (63%) and spine-necks (27%), and they also innervated interneuron dendrites (5%), while some (6%) postsynaptic targets could not be classified (Figure 1 O, Table S4). All innervated spines received another, putatively glutamatergic asymmetric, type I input from an unlabeled terminal, suggesting that, contrary to previous suggestion (*2*), cholinergic synapses alone do not induce spine formation. These data suggest that about 90% of cholinergic synapses target pyramidal cells in CA1, whereas they also innervate interneurons (at least 5%), which ratio is close to the neuronal ratios in CA1.

In somatosensory cortex S1, differentiation of pyramidal and interneuronal dendritic shafts is not possible based on electron microscopic profiles. In S1 cortex, cholinergic terminals targeted dendrites (54%), spines (38%), while 8% of their synaptic targets remained unidentified (Table S4).

#### Cholinergic synapses express GABA_A_ receptors and its scaffolding protein postsynaptically

Previously, we demonstrated that cholinergic synapses (in CA1, S1 somatosensory cortex, prefrontal cortex, basolateral amygdala and centrolateral thalamic nucleus) expressed the postsynaptic protein NL2 (*1*). This protein directly interacts with gephyrin, a core scaffolding protein of inhibitory postsynaptic densities (*3*) and their complex is implicated in the anchoring and clustering of GABA_A_ receptors postsynaptically (*4*–*6*). Here, we labeled cholinergic fibers either with vesicular acetylcholine transporter (vAChT, in WT mice) or eYFP (in ChAT-Cre mice, where medial septum was injected with a Cre-dependent eYFP-expressing AAV). Latter labelings were developed with DAB, while gephyrin or GABA_A_ receptor γ2 subunits were labeled with immunogold particles. Using electron microscopy, we tested fully reconstructed synapses of vAChT or eYFP-AAV labeled terminals for the presence of immunogold particles. Data from vAChT and AAV-eYFP labeled samples were not statistically different; therefore, they were pooled.

We found that at least 81% of 188 synapses collected in the CA1 area of two WT and two ChAT-Cre mice contained gephyrin postsynaptically (Table S3). Immunogold particles in these synapses were associated with the cytoplasmic side of the postsynaptic membrane of the innervated pyramidal cell dendrites (Figure 1B-D, G-I) and spines (Figure S1B) or interneuron dendrites (Figure 1J). The antibody used for GABA_A_ receptor-labeling was directed against an extracellular epitope of the γ2 subunit (*7*, *8*) and labeled cholinergic synaptic clefts accordingly. In the CA1 of three WT and two ChAT-Cre mice, at least 80% of the cholinergic synapses (out of 172) showed GABA_A_ receptor γ2 subunit labeling (Table S3). Cholinergic fibers established GABA_A_ receptor containing synapses on both dendrites (Figure 1E,F,K,L,N; Figure S1 C, D) and spine-necks (Figure 1M). In addition, we found that at least 83% of synapses of vAChT-positive terminals in the somatosensory cortex S1 (n=36, Figure S1 E-F) contained these GABA_A_ receptor γ2 subunits. Because reliable nicotinic receptor antibodies are not available, they could not be localized directly.

#### Cholinergic terminals possess the molecular machinery for GABA-release

By crossing a zsGreen fluorescent reporter-mouse-line with a vesicular GABA transporter (vGAT)-Cre mouse line, we created mice, in which all GABAergic cells are zsGreen labeled (vGAT-zsGreen mice). After co-labeling medial septum sections for ChAT, we found that all cholinergic cells were also positive for zsGreen (n=243 cells in 2 mice, Figure 2A, B), while many of the cells were positive only for zsGreen, corresponding to the non-cholinergic, GABAergic neurons of the medial septum. These results confirmed that all hippocampal projecting cholinergic cells express vGAT.

To confirm that septo-hippocampal cholinergic fibers can synthetize GABA, we performed immunofluorescent reactions against glutamate-decarboxylase (GAD65) and eYFP on hippocampal sections of ChAT-Cre mice, in which medial septal cholinergic fibers were labeled with Cre-dependent eYFP-AAV. We confirmed the GAD65 expression in eYFP positive terminals (Figure 2C) as well.

To confirm vGAT protein expression in cholinergic terminals we performed vGAT-eYFP and vAChT-eYFP multiple labelings, and found that vGAT was present in the majority of the eYFP positive cholinergic terminals (at least 82.9%, n=311 in 3 mice Figure 2D). We also quantified vAChT expression, which was found to be present in at least 63.6% of the eYFP positive cholinergic terminals (n=364 in 3 mice).

Using postembedding GABA-immunogold staining, we also tested, whether cholinergic terminals contain GABA itself (Figure 2 E-F). We measured postembedding GABA-immunogold labeling densities in preembedding labeled vAChT-positive terminals (n=2 mice, 24 terminals). Background level was estimated measuring gold particles on vAChT-negative, putative glutamatergic terminals that formed asymmetric synapses in the vicinity of the examined vAChT-positive terminals (n=2 mice, 34 terminals). Data from two mice were not different statistically; therefore, they were pooled. We found a significantly higher level of immunogold labeling for GABA in vAChT-positive terminals than in glutamatergic terminals (3.3 times higher density; Figure 2F), suggesting the presence of GABA in these terminals.

#### Synaptic vesicles of cholinergic terminals are highly heterogeneous and relatively large

We analyzed the volume and elongation of vesicles (*9*, *10*) in cholinergic terminals along with similar data from purely GABAergic terminals (Figure 4E). As expected, GABAergic vesicles were small and elongated (median volume: 13730 nm^3^, 11174-16434 nm^3^ interquartile range; median of elongation factor: 2.90, 2.52-3.41 interquartile range; n=54 vesicles from 2 mice). However, the volume of cholinergic vesicles were significantly larger (Mann-Whitney test, p<0.001). The volume of the vesicles in cholinergic terminals showed a significantly higher variability as well (F-test, p<0.001), ranging from the very large and round ones to the small and elongated vesicles (median volume: 23267 nm^3^, 19160-27539 nm^3^ interquartile range; median elongation factor: 1.80, 1.57-2.04 interquartile range; n=140 vesicles from two mice, Figure 4E). The small and elongated vesicles in the cholinergic terminals were similar to the purely GABAergic ones from GABAergic interneuron terminals (Figure 4E). These data suggest that cholinergic terminals contain both smaller, more elongated GABAergic vesicles and larger, rounder cholinergic vesicles, both of which are released from the same synaptic active zone. Interestingly, some vesicles that were directly labeled with vAChT-immunogold particles were rather large and round (Figure 4E). We also observed a vAChT-labeled vesicle, fused to the synaptic membrane, likely releasing its transmitter into the synaptic cleft (Figure 4C). In subsequent experiments, we collected further evidence that acetylcholine and GABA are filled into different vesicles allowing their separately-regulated co-transmission.

#### Acetylcholine and GABA are released at the same active zone in cholinergic terminals

We also tested, whether the two transmitter systems use the same or distinct active zones. We performed multiple immunofluorescent labeling experiments on virus labeled (eYFP) cholinergic fibers in ChAT-Cre mice for gephyrin, vAChT and eYFP, followed by confocal fluorescent imaging. We observed gephyrin-labeled puncta opposed to eYFP-positive terminals (Figure 4G, H), identifying the postsynaptic active zones of these fibers. vAChT-labeling was clearly concentrated opposite to the gephyrin-puncta, proving a tight association to the synaptic active zones (Figure 4G, H). Scale-free analysis confirmed that the likelihood of vAChT labeling was the highest at the synaptic active zones (Figure 4I; n=32 synapses from two mice). To directly examine the existence of a mixed cholinergic/GABAergic vesicle pool, we labeled brain slices for vGAT, vAChT and eYFP, and performed correlated fluorescent confocal laser scanning microscopy (CLSM) and superresolution STORM imaging (Figure 4J). The superresolution images confirmed that vAChT- and vGAT-labeled vesicle pools overlap, and were localized to the same small, confined portions of the eYFP septo-hippocampal terminals.

#### Acetylcholine and GABA are released from different vesicles in cholinergic terminals

Although a previous study in rat has suggested that GABA-containing synaptic vesicles do not contain acetylcholine (*11*), using a highly specific method, we confirmed that GABA and acetylcholine vesicular transporters are localized on different vesicles in mouse cortical axon terminals. We used isolated synaptic vesicles to test whether acetylcholine and GABA are packed into the same vesicles. Isolation from neocortex and hippocampus was performed according to Mutch et al. (Figure S4A, *12*). Isolated vesicles were investigated by flow cytometry for synaptophysin (SYP) expression. Labeling with a specific SYP antibody resulted in an about two orders of magnitude higher mean fluorescent intensity of vesicle preparations compared to the labeling with the secondary antibody alone (Figure S4B), suggesting a highly purified preparation. After fixation and dehydration of vesicle preparations, we confirmed the presence of synaptic vesicles surrounded by lipid bilayer on electron microscopic images (Figure S4E). The analysis confirmed that the diameter of the isolated vesicles was 37.55 nm (median, 33.78-40.21 nm interquartile range, n=100 vesicles; Figure S4C), in accordance with literature data (*13*). Next, we performed immunolabeling experiments on isolated synaptic vesicles fixed onto coverslips. Prior to CLSM imaging, we labeled the samples for SYP, vGAT, vAChT and vesicular glutamate transporter (VG1). As expected, we observed well separated fluorescent dots (point-spread functions, PSF) of the fluorophores in one single focal plane (Figure S4F), but most PSFs showed vesicular co-localization of one of the vesicular transporters and SYP. Control experiments of the immunolabeling confirmed the lack of unspecific staining (Figure S4I). In the absence of vesicle suspension, no PSFs were found in the CLSM scans, and the exclusion of any primary antibody led to the selective disappearance of PSFs in the corresponding channel. We also tested the distribution of fluorescent PSFs on the CLSM images. SYP-labeled vesicles were usually more than 1 μm away from each other as nearest-neighbor analysis of PSF centroids confirmed (median: 1.16 μm, 0.82-1.59 μm interquartile range, 0.46-3.92 min-max, n=149 vesicles; Figure S4D). When PSFs in different channels colocalized, their centroids were never farther away from each other than 0.130 μm (median: 0.03 μm, 10-50 μm interquartile range, 0-0.13 min-max, n=92 vesicles; Figure S4D). These experiments confirmed that co-localizing PSFs correspond to a single vesicle. Next, we analyzed co-localizations of the PSFs in different channels (Figure S4G) and found that 29.2% of vesicles were labeled only for SYP, 44.5% were double-labeled for VG1 and SYP, 14.3% were double-labeled for vGAT and SYP, 11.1% were double-labeled for vAChT and SYP. Only a negligible amount of vesicles (0.9%) were triple labeled with any combinations, whereas only a subfraction of these vesicles (0.14% of all) were co-labeled for vAChT and vGAT. Only 0.98% of all vGAT/SYP positive vesicles were labeled for vAChT, and only 1.26% of all vAChT/SYP positive vesicles were labeled for vGAT. These numbers are in the range of false positive labeling as confirmed in the control experiments, where primary antibodies were omitted (Figure S4I). These data suggest that vesicular transporters for glutamate, GABA and acetylcholine are expressed by distinct vesicle populations in cortical samples (Figure S4H; n=353 vesicles). Therefore, acetylcholine and GABA may be released at the same active zones, but from different vesicles.

#### Identification of basal forebrain cholinergic fibers in the hippocampus: control experiments

We either used immunolabeling against the vesicular acetylcholine transporter (vAChT), or performed anti-eYFP staining on sections from ChAT-Cre mice, the medial septal areas of which have previously been injected with Cre-dependent eYFP-adeno associated virus (AAV). Both of these methods had to be verified for selectivity and specificity, thus we completed a comprehensive set of control experiments. The cholinergic innervation of the hippocampus is reported to originate exclusively from the basal forebrain. Although the presence of a local cholinergic cell population in the mouse hippocampus was reported to be an artifact (*14*) we also tested for it. We injected Cre-dependent eYFP-AAV into the hippocampi of ChAT-Cre mice (Figure S2C, inset), and stained hippocampal sections for eYFP, vAChT and vGAT (Figure S2E, F). We found a few eYFP positive cells in the hippocampus. They were extremely rare and resembled dentate gyrus granule cells and CA3 pyramidal cells. We also found some sparsely distributed eYFP positive fibers originating from them, but vAChT or vGAT immunoreactivity was never found in these eYFP positive terminals (0 out of 323 terminals, from 2 mice, Figure S2F). We also tested vAChT positive terminals in the same samples, and never found any eYFP-positivity in them (0 out of 3673 from 2 mice, Figure S2F). Thus, we confirmed that there are no cholinergic cells in the hippocampus, only some extremely rare ectopic expression of the Cre enzyme. These results also confirmed that we can reliably label the septo-hippocampal cholinergic fibers with vAChT labeling.

To verify the other approach, we injected eYFP-AAV into the medial septal areas of ChAT-Cre mice (Figure S2C), and peformed PV/ChAT/eYFP triple labelings (Figure S2D). 97.6% of all tested eYFP positive cells in the MS were also positive for ChAT (the few % of false negative cells are likely due to not perfectly efficient antibody penetration), but none of them were positive for PV (n=212 in 2 mice). We also tested the fibers of these cells in the hippocampus, and performed a PV/vAChT/eYFP triple labeling (Figure S2A, B). We found that eYFP positive terminals colocalized with vAChT-labeling, but were never positive for PV (n=252 terminals from 2 mice). These results confirmed that eYFP positive fibers in these animals originate exclusively from cholinergic cells.

### SUPPLEMENTARY STATISTICAL DATA

#### Statistical details for Figures

**Figure 2F:** Medians (columns) and interquartile ranges (bars) of immunogold densities of GABA labeling in glutamatergic (Glut, median: 3.5 gold particles/ μm2, interquartile ranges: 1.5-5.3) and in VAChT-positive terminals (VAChT, median: 11.5 gold particles/ μm2, interquartile ranges: 6.8-22.7). Asterisk indicates significant difference (Mann-Whitney Test: p< 0.05). vAChT-negative terminals forming type I synapses were considered to be glutamatergic.

**Figure 3H:** Amplitude, 20-80% rise time and decay time of unitary GABAergic IPSCs from pyramidal cells (n=5) and inhibitory neurons (n=16). Box plots represent median values, with interquartile ranges, whiskers represent min/max values. Amplitude in pA: PCs: 37.28 (20.94, 61.61); INs: 61.56(46.78, 98.39), Mann-Whitney Test: p<0.05. Rise time (in ms) in PCs: 2.06 (1.62, 2.29), INs: 1.29 (1.12, 1.83); Mann-Whitney Test: not significant. Decay time (in ms): PCs: 16.29 (15.72, 25.94), INs: 11.35 (8.68, 14.10), Mann-Whitney Test: p<0.05.

**Figure 3I:** Averages of IPSC amplitudes for the 5 pulses presented on panel G show strong short-term depression (STD) of GABAergic transmission evoked by stimulating cholinergic fibers. **PCs:** 2 Hz: 1^st^ –42.89 (±21.79); 2^nd^ –25.74 (±13.7); 3^rd^ –25.63 (±14.85); 4^th^ –24.65 (±18.64); 5^th^ –21.54 (±17.04). 5 Hz: 1^st^ – 40.12(±18.32); 2^nd^ – 22.96(±12.17); 3^rd^ –18.68 (±10.9); 4^th^ – 20.45 (±14.9); 5^th^ – 19.36 (±17.28). 10 Hz: 1^st^ – 38.59(±13.62); 2^nd^ – 19.04(±12.67); 3^rd^ – 13.65(±8.57); 4^th^ – 12.44(±7.5); 5^th^ – 11.46(±7.44). 20 Hz: 1^st^ –37.10 (±24.18); 2^nd^ –11.98 (±12.24); 3^rd^ –9.82 (±10.35); 4^th^ – 9.13(±8.92); 5^th^ – 6.78 (±8.06). **INs**: 2 Hz: 1^st^ – 66.72(±18.33); 2^nd^ – 51.53(±19.44); 3^rd^ – 43.19(±15.35); 4^th^ – 41.44(±13.34); 5^th^ – 36.78(±13.36). 5 Hz: 1^st^ – 70.55(±18.31); 2^nd^ – 51.73(±16.27); 3^rd^ – 39.02(±15.12); 4^th^ – 32.58(±15.49); 5^th^ – 32.76(±13.22). 10 Hz: 1^st^ – 70.01(±14.75); 2^nd^ – 50.89(±14.89); 3^rd^ – 32.75(±14.93); 4^th^ – 29.50(±13.82); 5^th^ – 30.02(±14.41). 20 Hz: 1^st^ – 67.1(±15.22); 2^nd^ –35.0 (±15.22); 3^rd^ –28.76 (±15.07); 4^th^ – 23.68(±14.38); 5^th^ – 21.55(±13.14).

**Figure 5B:** IPSP amplitude at control (median (interquartile range)): 0.88 mV (0.78-1.29), atropine: 1.35 mV (1.13-1.38); Wilcoxon-sign rank test: p<0.05. EPSP amplitude at control: 0.42 mV (0.27-0.77), atropine: 0.65 mV (0.45-2.22); Wilcoxon-sign rank test: p<0.05.

**Figure 5C:** IPSP amplitude at control: 0.74 mV (0.43-1.05), AFDX-116: 1.08 mV (0.58-1.42); Wilcoxon-sign rank test: p<0.05; EPSP amplitude at control: 0.17 mV (0.1-0.27), AFDX-116: 0.29 mV (0.18-0.74); Wilcoxon-sign rank test: p<0.05.

**Figure 5D:** IPSP amplitude at control: 0.79 mV (0.57-1.05), CGP: 0.9 mV (0.79-1.24); Wilcoxon sign rank test: p<0.05; EPSP amplitude at control: 0.24 mV (0.13-0.25), CGP: 0.37 mV (0.23-0.46); Wilcoxon sign rank test: p<0.05.

**Figure 5E:** IPSP amplitude at control: 1.21 mV (1.01-1.56), ω-agatoxin: 0.85 mV (0.59-0.96); Wilcoxon sign rank test: p<0.05. EPSP integral at control: 0.29 mV*s (0.24-0.94); agatoxin: 0.34 mV*s (0.21-0.64), Wilcoxon sign rank test: p=0.63.

**Figure 5F:** EPSP integral at control: 0.55 mV*s (0.25-0.59); conotoxin: 0.13 mV*s (0.11-0.25); Wilcoxon sign rank test: p<0.05. IPSP amplitude at control: 0.74 mV (0.49-0.91), conotoxin: 0.68 mV (0.51-0.72); Wilcoxon sign rank test: p=0.52.

**Figure 5H:** EPSP integral at 2 Hz: 2.19 mV*s (0.41-3.63), 5 Hz: 2.56 mV*s (0.49-3.70), 10 Hz: 2.92 mV*s (0.53-3.88), 20 Hz: 1.82 mV*s (0.46-3.92); Wilcoxon-sign rank test: p<0.05; 0.05; 0.03.

**Figure 6D:** SWR rate in the absence of cholinergic blockers: 1.65 Hz (1.18-1.78) at control, 1.18 Hz (0.99-1.27) during illumination and 1.78 Hz (1.27-1.95) during recovery period, Wilcoxon-sign rank test: control-stimulation, p<0.05; stimulation-recovery, p<0.01; control-recovery, p<0.05.

**Figure 6F:** SWR rate in cholinergic blockers: 1.0 Hz (0.93-1.34) at control, 0.44 Hz (0.40-0.99) during illumination, 0.95 Hz (0.79-1.33) during recovery period, Wilcoxon-sign rank test: control-stimulation, p<0.01; stimulation-recovery, p<0.01; control-recovery, not significant.

**Figure 6I:** Epileptic discharge rate in the absence of AChR blockers: 0.61 Hz (0.47-0.68) at control, 0.54 Hz (0.44-0.58) during illumination, 0.62 Hz (0.5-0.63) during recovery period, Wilcoxon-sign rank test: control-stimulation, p<0.05; stimulation-recovery, p<0.05; control-recovery, not significant.

**Figure 6K:** Epileptic discharge rate in the presence of AChR blockers: 0.56 Hz (0.49-0.60) at control, 0.43 Hz (0.37-0.44) during illumination, 0.48 Hz (0.42-0.54) during recovery period, Wilcoxon-sign rank test: control-stimulation, p<0.01; stimulation-recovery, p<0.05; control-recovery, not significant.

### SUPPLEMENTARY DISCUSSION

#### Cholinergic non-synaptic neurotransmission

For decades, the predominant form of cholinergic communication was thought to be a form of “non-synaptic volume transmission" (*15*–*21*), which was supported by electron microscopic studies showing that cholinergic terminals form few synapses [3-17%: in cat striate cortex (*22*), rat parietal cortex (*23*, *24*), rat hippocampus (*25*, *26*), mouse hippocampus (*27*)], while some papers reported more frequent synapses [44-67%: in macaque prefrontal cortex (*28*), human temporal lobe (*29*), rat parietal cortex (*30*)]. Although acetylcholine esterase (AChE) was known to be highly effective in terminating extracellular cholinergic signal, the presence of certain extrasynaptic acetylcholine receptors (”receptor mismatch“, *27*, *31*) suggested that extracellular diffusion of acetylcholine occurs. Micro-dialysis experiments (*32*) and the localization of AChE, distant from cholinergic terminals, also seemed to support a non-synaptic “volume” transmission hypothesis (*15*). However, later, highly sensitive microelectrodes showed faster, phasic changes in extracellular acetylcholine levels that facilitated cue detection and cortical information processing (*21*, *33*–*36*). Basal forebrain cholinergic neurons were also shown to respond to reward and punishment with extremely high speed and precision (*37*), and recent data suggested that cholinergic cells may regulate cortical information processing with a remarkable, millisecond-scale temporal precision (*35*, *38*–*40*). However, such a delicate temporal precision is hard to imagine without synapses and it remains inconclusive, whether the mode of acetylcholine signaling is synaptic “wired” transmission or “non-synaptic volume” transmission by ambient acetylcholine (*27*).

#### GABAergic markers in cholinergic cells

Basal forebrain cholinergic cells share a common developmental origin with different populations of cortical, striatal and basal forebrain GABAergic neurons (*41*–*44*). Previous studies have suggested that less than 2% of cholinergic cells express GABAergic markers (*45*, *46*), while about 8% of ChAT-positive boutons in the cat striate cortex was shown to contain GABA (*47*). The recognition of GABAergic signaling in the BF cholinergic system may have been hampered by its lack of GAD67 (*48*) and GABA transporter 1 (*49*). Although the associations of cholinergic terminals with gephyrin (*50*) and NL2 (*1*) suggested the capability of GABAergic signaling from these terminals. While, for example, GABA is released together with glutamate or aspartate in the hippocampus, or with dopamine in periglomerular cells (*51*, *52*), and acetylcholine is released with glutamate in striatum (*53*); GABA and acetylcholine were also shown to be released together in retina and frontal cortex (*46*, *48*, *54*, *55*). However, the precise architecture, the mechanism of the dual cholinergic/GABAergic transmission and their hippocampal synaptic physiological and network effects have not yet been investigated.

### SUPPLEMENTARY EXPERIMENTAL PROCEDURES

#### Animals and surgery

A total of 21 (25-80 days old) male C57BL/6J mice, 51 (30-200 days old) ChAT-IRES-Cre mice from either sex (Jackson Laboratory, RRID: IMSR_JAX:006410, postnatal day 30–200) and 4 (30-50 days old) VGAT-IRES-Cre/Gt(ROSA)26Sor_CAG/ZsGreen1 mice were used in the present study. All experiments were performed in accordance with the Institutional Ethical Codex, Hungarian Act of Animal Care and Experimentation (1998, XXVIII, section 243/1998) and the European Union guidelines (directive 2010/63/EU), and with the approval of the Institutional Animal Care and Use Committee of the Institute of Experimental Medicine of the Hungarian Academy of Sciences. All efforts were made to minimize potential pain or suffering and to reduce the number of animals used.

ChAT-Cre mice were anesthetized with isoflurane followed by an intraperitoneal injection of an anesthetic mixture (containing 8.3 mg/ml ketamine, 1.7 mg/ml xylazine-hydrochloride in 0.9 % saline, 10 ml/kg bodyweight) and then were mounted in a stereotaxic frame. For selective labeling of septo-hippocampal cholinergic axons, we injected 30-60 nl AAV2/5-EF1a-DIO-eYFP (UNC Vector Core) or AAV2/5-EF1a-DIO-ChR2(H134R)-eYFP-WPRE-hGH (plasmid: Addgene 20298, Penn Vector Core) into the medial septal area. The coordinates for the injection were based on the atlas by Paxinos, G and Franklin, (2012): 1.0 mm anterior from the bregma, in the midline, and 4.3 mm below the level of the horizontal plane defined by the bregma and the lambda (zero level). For a control experiment, 100 nl injections of the same virus were delivered into the hippocampus (both hemispheres) of a ChAT-Cre mouse (coordinates: -2.7 mm from the bregma, +/- 2.5 mm from the midline, 2.5 mm from the zero level and -3.1 from the bregma, +/- 3 mm from the midline and 3 mm from the zero level). For the injections, we used a Nanoject 2010 precision microinjector pump (WPI, Sarasota, FL 34240). We used borosilicate micropipettes (Drummond, Broomall, PA) for the injections with tips broken to 40-50 μm. After the surgeries, the animals received 0.5–0.7 ml saline for rehydration and 0.03–0.05 mg/kg meloxicam as nonsteroidal anti-inflammatory drug (Metacam, Boehringer Ingelheim, Germany) intraperitoneally to support recovery, and we placed them into separate cages to survive for at least three weeks before decapitations or perfusions.

#### Slice preparation and recording conditions

After 3-6 weeks following injection (to reach an appropriate expression level in long-range projecting axons) horizontal slices were prepared. In all cases, mice were decapitated under deep isoflurane anesthesia. The brain was removed into ice-cold cutting solution, which had been bubbled with 95% O_2_ -5% CO_2_ (carbogen gas) for at least 15 min before use. The cutting solution contained the following (in mM): 205 sucrose, 2.5 KCl, 26 NaHCO_3_, 0.5 CaCl_2_, 5 MgCl_2_, 1.25 NaH_2_PO_4_, 10 glucose, saturated with 95% O_2_-5% CO_2_. Horizontal hippocampal slices of 300 μm or 450 μm thicknesses (for whole-cell and LFP recordings, respectively) were cut using a vibratome (Leica VT1000S). After acute slice preparation the slices were placed into an interface-type holding chamber for recovery. This chamber contained standard artificial cerebrospinal fluid (ACSF) at 35°C that gradually cooled down to room temperature. The ACSF had the following composition (in mM): 126 NaCl, 2.5 KCl, 26 NaHCO_3_, 2 CaCl_2_, 2 MgCl_2_, 1.25 NaH_2_PO_4_, 10 glucose, saturated with 95% O_2_-5% CO_2_. After incubation for at least 1.5 h, slices were transferred individually into a submerged-style recording chamber equipped with a single superfusion system. In case of LFP recording of hippocampal network activities, a double superfusion system was used for improved slices maintenance conditions (*57*, *58*). In the latter design, the slices were placed on a metal mesh and two separate fluid inlets allowed ACSF to flow both above and below the slices with a rate of 3–5 ml/min for each flow channel at 30–32°C (Supertech Instruments; www.super-tech.eu). The composition of the modified ACSF (mACSF) used in experiment presented in Figure 6 G-K was the following (in mM): 126 NaCl, 3.5 KCl, 26 NaHCO_3_, 1.6 CaCl_2_, 1.2 MgCl_2_, 1.25 NaH_2_PO_4_, 10 glucose saturated with 95% O_2_-5% CO_2_. Epileptic events were evoked by further increasing the potassium concentration from 3.5 to 8.5 mM after 15 minutes spent in mACSF in the recording chamber. Slices were visualized with an upright microscope (Nikon Eclipse FN1 or Olympus BX61WI) with infrared-differential interference contrast optics. Standard patch electrodes were used in all recording configurations (i.e., in whole-cell patch-clamp and field potential recordings). Pipette resistances were 3–6 MΩ when filled either with the intracellular solution or with ACSF. ACSF filled pipettes placed into CA1 pyramidal layer were used for local field potential (LFP) recordings. The composition of the intracellular pipette solution was the following (in mM): 110 K-gluconate, 4 NaCl, 20 HEPES, 0.1 EGTA, 10 phosphocreatine, 2 ATP, 0.3 GTP, 3 mg/ml biocytin adjusted to pH 7.3–7.35 using KOH (285–295 mOsm/L). Where indicated, high-chloride containing intracellular solution was used: 54 K-gluconate, 4 NaCl, 56 KCl, 20 HEPES, 0.1 EGTA, 10 phosphocreatine, 2 ATP, 0.3 GTP, 3 mg/ml biocytin adjusted to pH 7.3–7.35 using KOH (285–295 mOsm/L). Whole-cell series resistance was in the range of 5–15 MΩ. Series resistance was not compensated but was frequently monitored, and cells where the values changed more than 25% during recording were discarded from further analysis. Voltage measurements were not corrected for the liquid junction potential.

#### Drugs

To avoid polysynaptic effects in response to optical stimulation of cholinergic fibers, glutamatergic excitatory currents were blocked by 20 μM NBQX and 50 μM AP-5 in all experiments presented on Figure 3, Figure S3 and Figure 5. The following drug concentrations were used in the specified experiments (wash application): 20 μM NBQX, 50 μM AP5, 10 μM atropine, 1 μM MLA, 1 μM DHBE, 10 μM gabazine, 1 μM CGP-55845. For puff application of ω-conotoxin (1 μM) and ω-agatoxin (1 μM), a glass capillary was placed adjacent to the recorded cell. These drugs were injected by mouth-applied pressure for 1-2 minutes during the stimulation protocol following the control period. Drugs were dissolved in HEPES-based buffer with a composition in mM: 126 NaCl, 10 glucose, 2.5 KCl, 1.25 NaH_2_PO_4_, 2 CaCl_2_, 2 MgCl_2_, 26 HEPES, pH 7.3. Gabazine (SR 95531 hydrobromide), CGP-55845, MLA (Methyllycaconitine citrate), DhβE (Dihydro-β-erythroidine hydrobromide) and AF-DX 116 were purchased from Tocris Bioscience, ω-conotoxin and ω-agatoxin were purchased from Almone labs, and NBQX (2,3-Dioxo-6-nitro-1,2,3,4-tetrahydrobenzo[f]quinoxaline-7-sulfonamide) and AP5 were purchased from Hello Bio. All other salts and drugs were obtained from Sigma-Aldrich or Molar Chemicals KFT.

#### Anatomical identification

The recorded cells were filled with biocytin. After the recording, the slices were fixed in 4% paraformaldehyde in 0.1 M phosphate buffer (PB; pH 7.4) for at least 3 h, followed by washout with PB several times. Then, sections were blocked with normal goat serum (NGS; 10%) diluted in Tris-buffered saline (pH 7.4), followed by incubations in Alexa Fluor 594-conjugated streptavidin (1:1000; Invitrogen). Sections were then mounted on slides in Vectashield (Vector Laboratories) and were morphologically identified on the basis of their location, dendritic and axonal arborization.

#### Optogenetic illumination

For illumination, we used a blue laser diode (447 nm, Roithner LaserTechnik GmbH) attached to a single optic fiber (Thorlabs) positioned above the hippocampal slice. In all cases, 1 ms pulse length was used. In case of structured illumination (Figure S3), the same laser was used with a digital micromirror device (Polygon400, Mightex Systems, Toronto, Canada) integrated into the optical path of the microscope. We have used Polygon400 for the illumination of the slice at a remote location (~5-600 μm), to avoid direct illumination of ChR2-expressing terminals on the recorded cell.

#### Electrophysiological recordings and analysis

Both whole-cell and LFP recordings were performed with a Multiclamp 700A or 700B amplifier (Molecular Devices) and were low-pass filtered at 3 kHz using the built-in Bessel filter of the amplifier. Data were digitized at 10 kHz with a PCI-6042E board (National Instruments) and recorded with EVAN 1.3 software (courtesy of Prof. I. Mody, University of California Los Angeles, Los Angeles, CA). All data were analyzed off-line using custom-made programs written in MATLAB 8.5.0, Python 2.7 by D.S., and Delphi 6.0 by A.I.G.

The kinetic properties (20–80% rise times, decay time constant, latency) were calculated on stimulus triggered averaged events. Fitting of a single exponential function to the decaying phase of averaged responses and statistical analyses were performed in OriginPro, version 9.2.214 (OriginLab Corporation, Northampton, MA, USA). Latency from stimulation start to PSP onset was calculated as the time when the signal crossed 3 times standard deviation of baseline. Transmission probability was defined, as the probability of detectable IPSC in response to the optical stimulation. For characterizing short-term plasticity, each stimulation frequency was repeated 20 times with an interval of 3 seconds. To record in parallel the changes in cholinergic EPSP and GABAergic IPSPs (for the drug applications presented on Figure 5) cells were measured at -50 mV in current clamp mode. Since the amplitude of both PSPs increased, qualitative changes in amplitude could be measured, however in this way, we may have underestimated the changes in amplitude due to conductances overlapping in time and acting in the opposite directions. In the case of VDCC blocker application, we used five pulses to evoke larger cholinergic responses. We quantified EPSP area changes for ω-agatoxin and ω-conotoxin application after low-pass filtering (at 2 Hz) the response (e.g. subtracting GABAergic component).

Figure 5 data were normalized for visualization (all values were divided by the mean of the control sample (*59*). LFP signals were filtered with a two way RC filter to preserve phase. SWRs were pre-detected on 30 Hz low-pass-filtered field recordings using a threshold value of four times the SD of the signal. The pre-detected SWRs were then redetected using a program that detected SWR peaks and eliminated recording artifacts similar to SWRs. Epileptic event peaks were detected similarly. Event rates were calculated with equal time periods for control, stimulation and recovery phases. All automatic detection steps were manually supervised.

#### Perfusion

For perfusion, mice were deeply anesthetized as above. Mice for electron microscopical reconstruction of axonal segments and immunofluorescent labelings were perfused transcardially first with 0.9 % NaCl in 0.1M phosphate buffer (PBS) solution for 2 min followed by 4% paraformaldehyde (PFA) in 0.1 M phosphate buffer (pH=7.4; PB) for 40 min and finally with PBS for 10 min. In the case of one WT mouse for reconstruction the fixative also contained 0.5% glutaraldehyde. Two WT mice for postembedding GABA-labeling and electron tomography were perfused first with PBS for 2 min followed by a fixative containing 2% PFA and 2% glutaraldehyde in 0.1 M sodium acetate buffer (pH 6.5) for 2 min, and then by 2% PFA/2% glutaraldehyde in 0.1 M sodium borate buffer (pH 8.5) for 1 h (*60*). Brains were removed from the skull and postfixed overnight at 4°C in 2% PFA/2% glutaraldehyde in 0.1 M sodium borate buffer (pH 8.5). Mice for electron microscopical or immunfluorescent detection of gephyrin and GABA_A_ receptors were perfused transcardially with ice-cold, oxygenated artificial cerebrospinal fluid [containing (mM) NaCl 125, KCl 2.5, CaCl_2_ 2.5, MgCl_2_ 2, NaHCO_3_ 26, NaH_2_PO_4_ 1.25, glucose 25; Notter et al., 2014]. Then the brain was removed from the skull and cut in blocks containing the hippocampal formation (separated hemispheres) and the medial septal area. The blocks were incubated in a fixative containing 4% PFA and 0.2% glutaraldehyde in PB for 90 min at 4C and then embedded in 2% agarose for sectioning. Coronal sections were cut on a Leica VT1200S vibratome at 50 μm. All series of sections were rinsed in PB, cryoprotected sequentially in PB containing 10% and 30% sucrose, frozen in liquid nitrogen and stored at –70°C until further processing.

#### Antibodies

We have summarized the primary antibodies used, their concentrations and information on their specificity in Table S1. The secondary antibodies (see Table S2) were extensively tested for possible cross-reactivity with the other secondary or primary antibodies, and possible tissue labeling without primary antibodies was also tested to exclude autofluorescence or specific background labeling by the secondary antibodies. No specific-like staining was observed under these control conditions.

#### Single and double-labeling preembedding immunoelectron microscopy

Sections were freeze-thawed two times over liquid nitrogen and washed in PB. Sections for gephyrin immunolabeling were treated with 1% sodium borohydride in PB for 10 min. For detection of GABA_A_ receptors, the sections were incubated in 0.2 M HCl solution containing 2 mg/ml pepsin (Dako) at 37°C for 2-4 min. After extensive washes in PB and 0.05 M Tris-buffered saline (pH 7.4; TBS) sections were blocked in 1 % human serum albumin (HSA; Sigma-Aldrich) in TBS. Then, they were incubated in a mixture of primary antibodies (Table S1) diluted in TBS containing 0.05% sodium azide for 2-3 days. After repeated washes in TBS, the sections were incubated in blocking solution (Gel-BS) containing 0.2 % cold water fish skin gelatine and 0.5 % HSA in TBS for 1 h. Next, sections were incubated in gold-conjugated and biotinylated secondary antibodies (Table S1) diluted in Gel-BS overnight. After extensive washes in TBS the sections were treated with 2 % glutaraldehyde in 0.1 M PB for 15 min to fix the gold particles into the tissue. This was followed by incubation in avidin–biotinylated horseradish peroxidase complex (Elite ABC; 1:300; Vector Laboratories) diluted in TBS for 3 h at room temperature or overnight at 4°C. The immunoperoxidase reaction was developed using 3,3-diaminobenzidine (DAB; Sigma-Aldrich) as chromogen. To enlarge immunogold particles, sections were incubated in silver enhancement solution (SE-EM; Aurion) for 40-60 min at room temperature. The sections were then treated with 1% (for electron tomography) or 0.5 % OsO_4_ in 0.1 M PB, at room temperature (for electron tomography) or on ice, dehydrated in ascending alcohol series and in acetonitrile and embedded in Durcupan (ACM; Fluka). During dehydration, the sections were treated with 1% uranyl acetate in 70% ethanol for 20 min. For electron microscopic analysis, tissue samples from the CA1 area of dorsal hippocampus/ somatosensory cortex (S1) were glued onto Durcupan blocks. Consecutive 70 nm-thick (for conventional electron microscopic analysis) or 150 nm-thick (for electron tomography) sections were cut using an ultramicrotome (Leica EM UC6) and picked up on Formvar-coated single-slot grids. Ultrathin sections for conventional electron microscopic analysis were counterstained with lead citrate (Ultrostain 2, Leica) and examined in a Hitachi 7100 electron microscope equipped with a Veleta CCD camera (Olympus Soft Imaging Solutions, Germany). 150 nm-thick electron tomography sections were examined in FEI Tecnai Spirit G2 BioTwin TEM equipped with an Eagle 4k HS camera.

#### Combination of preembedding and postembedding immunocytochemistry for electron microscopy

VAChT was visualized using the preembedding gold method as described above. Alternate serial 70 nm-thick sections were mounted on copper and nickel grids (5-6 sections/ grid). Postembedding GABA-immunostaining was carried out on nickel grids according to a modified protocol (*62*). Incubations of sections were performed on droplets of solutions in humid Petri dishes in the following order: 0.5% periodic acid (H_5_IO_6_) for 5 min at room temperature; three times at 2 min wash in distilled water; 3 min in TBS; 15 min in 1% ovalbumin dissolved in TBS at 37°C; 8 min in TBS, 90 min rabbit anti-GABA antiserum (1:10 000 in TBS) at 37°C; two times at 10 min TBS; 10 min in TBS containing 1% BSA and 0.05% Tween 20; 90 min at room temperature in 10 nm colloidal gold-conjugated goat anti-rabbit IgG (BBI solutions; 1:1000 in the same solution as before); 3 times at 5 min wash in distilled water; 20 min in 10% saturated uranyl acetate; 4 wash in distilled water; staining with lead citrate; wash in distilled water. The etching procedure of postembedding GABA-immunostaining removes the silver precipitate of the preembedding vAChT-labeling (see Figure 2E); therefore, only every second electron microscope grid was reacted for GABA and the analysis of vACHT-positive terminals were carried out using the so called mirror technique. Sections on copper grids adjacent to the first sections on nickel grids were systemically scanned for preembedding immunogold-labeled vAChT-positive terminals. These terminals were identified in the next, GABA-labeled section, followed in consecutive serial sections and digital images were taken at 50,000 times magnification in each serial section. Postembedding immunogold particles were counted on these GABA-labeled terminals and the examined surface area was measured using the Reconstruct software (*63*). For comparison, we have also measured the immunogold densities of putative glutamatergic terminals using the same method, in the same series of images. Terminals forming type I (asymmetric) synapses were considered glutamatergic. Although periodic acid-etching of the sections is known to remove preembedding antibodies together with immunogold and silver precipitation, we confirmed this again in a control experiment because, here, both the preembedding labeling and the postembedding GABA staining was performed using rabbit antibodies. In these control experiments, the postembedding GABA immunostaining was carried out without the primary anti-GABA antibody, in which case practically no postembedding gold particles were detected, confirming the specificity of the method.

#### Electron microscopy analysis of cholinergic synapses

For evaluation of the gephyrin and GABA_A_ receptor content at synapses of cholinergic axons, we performed preembedding double labeling. VAChT (in wild type mice) or eYFP (in ChAT-Cre mice, see above) was labeled with DAB, while gephyrin or GABA_A_ γ2 subunit was labeled with immunogold (see above, Table S1). Electron microscopic serial sections were systemically scanned for synapses of DAB-labeled terminals. Parallel appositions between the membranes of a DAB-containing terminal and a putative postsynaptic target were regarded as a synapse if they displayed widening of the extracellular space at the presumptive synaptic cleft and clustered synaptic vesicles in the terminal. Synapses found were followed and photographed at 30,000 magnification in every section, where they were present throughout the series: thus they were fully reconstructed. Since the background labeling (measured in putative glutamatergic synapses in the same series of sections) was negligible, synapses containing at least one gold particle were regarded as positive.

#### 3D reconstruction of axonal segments

Cholinergic axonal segments (n=17) were reconstructed using consecutive serial 70 nm-thick sections double-labeled for choline acetyltransferase (ChAT; DAB) and neuroligin 2 (NL2; immunogold) in wild type mice (n=2) or eYFP (DAB) and gephyrin (immunogold) in ChAT-Cre animals, in which the medial septum was injected with Cre-dependent eGFP-containing virus (n=2). CB_1_-positive axons were reconstructed from a series of 70 nm-thick sections double labeled for CB_1_ (DAB) and NL2 (immunogold). DAB-containing axons were followed in consecutive serial sections and digital images were taken at 30,000 times magnification in each serial section. Three-dimensional reconstructions of plasma membranes, mitochondria, putative synapses of DAB-containing axons and postsynaptic gold particles were made using the Reconstruct software (*63*). For preparing figures of reconstructed axons and measuring their 3d length, Blender software (www.blender.org) was used. Postsynaptic targets of cholinergic axons were classified as described earlier (*64*). Briefly, spines were recognized by their small size and specific morphology. In CA1 str. oriens and radiatum, dendrites that have spines and do not receive type I (asymmetric) inputs on their shafts are known to be pyramidal cells (*65*), whereas dendrites receiving type I synapses on their shafts are interneurons (*66*). The robustness of this classification method was reconfirmed recently (*64*). Because in str. lacunosum-moleculare the shafts of pyramidal dendrites may receive type 1 inputs (*65*) they were not distinguished from interneuron dendritic shafts. Also, the dendritic shafts in somatosensory cortex S1 could not be identified based on morphological features.

#### Immunofluorescent labeling and confocal laser-scanning microscopy

Before the immunofluorescent staining, the 50 μm thick sections were washed in PB and Tris-buffered saline (TBS). This was followed by blocking for 1 hour in 1% human serum albumin (HSA) and 0.1% Triton X-100 dissolved in TBS. After this, sections were incubated in mixtures of primary antibodies overnight at room temperature. After incubation, sections were washed in TBS, and were incubated overnight at 4°C in the mixture of, all diluted in TBS. Secondary antibody incubation was followed by washes in TBS, PB, the sections were mounted on glass slides, and coverslipped with Aqua-Poly/Mount (Polysciences). Immunofluorescence was analyzed using a Nikon Eclipse Ti-E inverted microscope (Nikon Instruments Europe B.V., Amsterdam, The Netherlands), with a CFI Plan Apochromat VC 60XH oil immersion objective (numerical aperture: 1.4) and an A1R laser confocal system. We used 405, 488, 561 and 647 nm lasers (CVI Melles Griot), and scanning was done in line serial mode. Image stacks were obtained with NIS-Elements AR software, and deconvolved using Huygens Professional software (www.svi.nl).

#### STORM super-resolution microscopy

Before the immunofluorescent staining for the super-resolution experiments, the 20 μm thick sections were washed in PB and TBS. This was followed by blocking for 1 hour in 1% HSA dissolved in TBS. After this, sections were incubated in mixtures of primary antibodies; overnight at room temperature. After incubation, sections were washed in TBS, and sections were incubated overnight at 4°C in the mixtures of secondary antibodies. Secondary antibody incubation was followed by washing in TBS, PB; then hippocampi were cut out with scalpels in buffer, sections were dried on clean coverslips, and stored for maximum 3 weeks at room temperature in a dry environment before imaging. 3D direct-STORM (direct Stochastic Optical Reconstruction Microscopy) acquisition protocol was used as described before (*67*). The imaging setup was built around a Nikon Ti-E inverted microscope equipped with a Perfect Focus System, with an Andor iXon Ultra 897 EMCCD camera, a C2 confocal head, 405, 488 and 561 nm lasers (Melles Griot 56RCS/S2780, Coherent Sapphire), and a high power 647 nm laser (300 mW, MPB Communications VFL-P-300-647). A high NA 100x oil immersion objective (Nikon CFI SR Apochromat TIRF 100x oil, 1.49 NA) was used for imaging. During STORM acquisition the emitted light was let through 582/636 nm and 670/760 nm bandpass filters to reach the detector for the 561 and 647 channels, respectively. We used the NIS-Elements AR 4.3 with N-STORM 3.4 software for acquisition and analysis. Imaging medium was mixed from 80 μl DPBS (Dulbecco’s phosphate-buffered saline), 10 μl MEA (mercaptoethylamine) solution, 10 μl 50% glucose solution, and 1 μl GLOX (glucose oxidase) solution. Coverslips with the dried sections were mounted onto microscope slides with 25 μl of freshly prepared imaging medium, and sealed with nail-polish. After selecting the region of interest, confocal image stacks were acquired containing 15 focal planes with 80 x 80 x 150 nm voxel size in x, y and z, respectively. This was followed by bleaching the fluorophores in the STORM channels (561 and 647) with high intensity laser illumination, and running dSTORM acquisition using oblique (near-TIRF) illumination. The acquisition in the two channels was done in sequential mode. Confocal stacks were deconvolved (Huygens Professional), and – together with the STORM molecule lists – processed for further analysis in VividSTORM software (see "Analysis”). Imaging was performed with identical parameters (depth in the section, laser intensities, etc.) for all samples.

#### Electron tomography

For the electron tomographic investigation we used 150 nm thick sections from the hippocampal CA1 region from the anti-vAChT immunogold stained material (see: “Pre-embedding immunoelectron-microscopy”). Before electron tomography, serial sections on single-slot copper grids were photographed with a Hitachi H-7100 electron microscope and a Veleta CCD camera. Serial sections were examined at lower magnification, and vAChT-positive synaptic terminals from the CA1 area selected. For each terminal, the section containing the largest synaptic cross section was chosen for electron tomography. After this, grids were put on drops of 10% HSA in TBS for 10 minutes, dipped in distilled water (DW), put on drops of 10 nm gold conjugated Protein-A in DW (1:3), and washed in DW. Finally, we deposited 5 and 5 nm thick layers of carbon on both sides of the grids. Electron tomography was performed using a Tecnai T12 BioTwin electron microscope equipped with a computer-controlled precision stage (CompuStage, FEI). Acquisition was controlled via the Xplore3D software (FEI). Regions of interest were pre-illuminated for 4-6 minutes to prevent further shrinkage. Dual-axis tilt series were collected at 2 degree increment steps between -65 and +65 degrees at 120 kV acceleration voltage and 23000x magnification with -1.6 – -2 μm objective lens defocus. Reconstruction was performed using the IMOD software package (*68*). Isotropic voxel size was 0.49 nm in the reconstructed volumes. After combining the reconstructed tomograms from the two axes, the nonlinear anisotropic diffusion (NAD) filtering algorithm was applied to the volumes. Segmentation of the terminals has been performed on the virtual sections using the 3Dmod software, and measurements were done on the scaled 3D models.

#### Isolation of synaptic vesicles

Synaptic vesicles were isolated according to (*12*), with some modifications. Briefly, cortices and hippocampi of ten C57Bl6 mice (80 days old, mixed gender) were pulverized mechanically after freezing in liquid nitrogen. The resulting fine powder was then homogenized using a Teflon-glass homogenizer in 0.3M sucrose-containing HEPES buffer (0.3 M sucrose, 50 mM HEPES, pH 7.4, 2 mM EGTA). The homogenate was centrifuged at 100 000g (1h, 4°C). The supernatant was laid onto a 0.6M/1.5M sucrose step gradient and centrifuged at 260 000g (2h, 4°C). Synaptic vesicles were collected from the 0.6M/1.5M sucrose solution interface. The vesicles were either immediately used or were frozen in liquid nitrogen and stored at -20°C for later use.

#### Immunostaining and flow cytometry analysis of isolated synaptic vesicles

The vesicles were dialyzed overnight, at 4°C, in phosphate buffered saline (PBS; 800ml). For dialysis a 300kD MWCO membrane was used (Biotech CE Dialysis Tubing). All solutions used during the dialysis or subsequent immunostaining were thoroughly filtered through either 0.2μm (Millipore) or 0.02μm pore diameter filters (Anodisc-47; GE Healthcare/Whatman). The vesicles were incubated with primary antibody in the presence of 0.1% BSA (RT, 1 h) followed by incubation with secondary antibody, (RT, 1h), under constant shaking. Prior to FACS analysis the stained vesicle suspensions were dialyzed again against PBS (800 ml, at RT, for 1h). FACS analysis was performed on a BD FACSVerse instrument.

#### Transmission electron microscopy and confocal laser scanning microscopy of isolated synaptic vesicles

Isolated synaptic vesicle suspension was diluted 10x in TBS, and 100 μL was dropped onto ultra-clean coverslips. After 10 minutes, 100 μL 4% PFA in 0.1 M PB was added to the drops. For a further 15 minutes the coverslips were washed gently in distilled water (DW), and processed for TEM and CLSM analysis. For the electron microscopic investigation, vesicles fixed to the coverslips were treated with 1% OsO4 in 0.1 M PB at room temperature, dehydrated in ascending alcohol series and in acetonitrile, and embedded in Durcupan (ACM; Fluka). During dehydration, the coverslips were treated with 1% uranyl acetate in 70% ethanol for 20 min. After resin polymerization, small pieces were cut and removed from the coverslip surface, glued onto plastic blocks, and 40 nm thick sections were prepared using an ultramicrotome (Leica EM UC6) and picked up on Formvar-coated single-slot grids. Ultrathin sections were examined in a Hitachi 7100 electron microscope equipped with a Veleta CCD camera (Olympus Soft Imaging Solutions, Germany). For CLSM analysis coverslips were washed in PB and Tris-buffered saline (TBS). This was followed by blocking for 1 hour in 1% human serum albumin (HSA) dissolved in TBS. After this, sections were incubated in mixtures of primary antibodies for 1 hour at room temperature. After incubation, sections were washed in TBS, and were incubated for 1 hour at room temperature in the mixture of secondary antibodies, all diluted in TBS. Secondary antibody incubation was followed by washes in TBS, PB, DW and the coverslips were mounted on glass slides with Aqua-Poly/Mount (Polysciences). Immunofluorescence was analyzed using a Nikon Eclipse Ti-E inverted microscope (Nikon Instruments Europe B.V., Amsterdam, The Netherlands), with a CFI Plan Apochromat VC 60XH oil immersion objective (numerical aperture: 1.4) and an A1R laser confocal system. The possible influence of chromatic aberration on PSF-position was controlled using fluorescent TetraSpeck™ Microspheres (0.1 μm diameter, blue/green/orange/dark red, ThermoFisher Scientific). We used 405, 488, 561 and 647 nm lasers (CVI Melles Griot), and scanning was done in line serial mode. Image stacks were obtained with NIS-Elements AR software, and deconvolved using Huygens Professional software (www.svi.nl).

#### Statistical analysis

During segmentation of tomographic volumes, z-scaling was calculated from the thickness difference of the reconstructed volume and the original section thickness, and applied to the 3D models. Mesh surface areas and volumes inside meshed objects were measured with the “imodinfo” program.

In STORM imaging “molecule lists” were exported from NIS in txt format, and the 3 image planes of the ics-ids file pairs from the deconvolved confocal stacks matching the STORM-volume were converted to the ome-tiff format using Fiji software. Confocal and corresponding STORM images were fitted in VividSTORM (*69*).

When data populations in this work had a Gaussian distribution according to the Shapiro-Wilks W test, we reported parametric statistical features (mean ± SD), otherwise we reported non-parametric statistical features (median, interquartile range). For the presentation of electrophysiological data we used median and interquartile range, because the data did not show Gaussian distribution. Two non-parametric groups were compared using the Mann– Whitney U test. Wilcoxon signed-rank test was used for calculating significance between dependent non-parametric data groups. F-test was used to compare the variability of data groups. The types of statistical tests used in different investigations are indicated in the text. We always used two-tailed statistical tests. All statistical analyses were carried out using the software package Statistica (StatSoft, Tulsa, OK, USA) or OriginPro 9.2.214 (OriginLab Corporation, Northampton, MA, USA). The differences were considered significant at p<0.05.

### SUPPLEMENTARY FIGURES

**Supplementary Figure S1.**
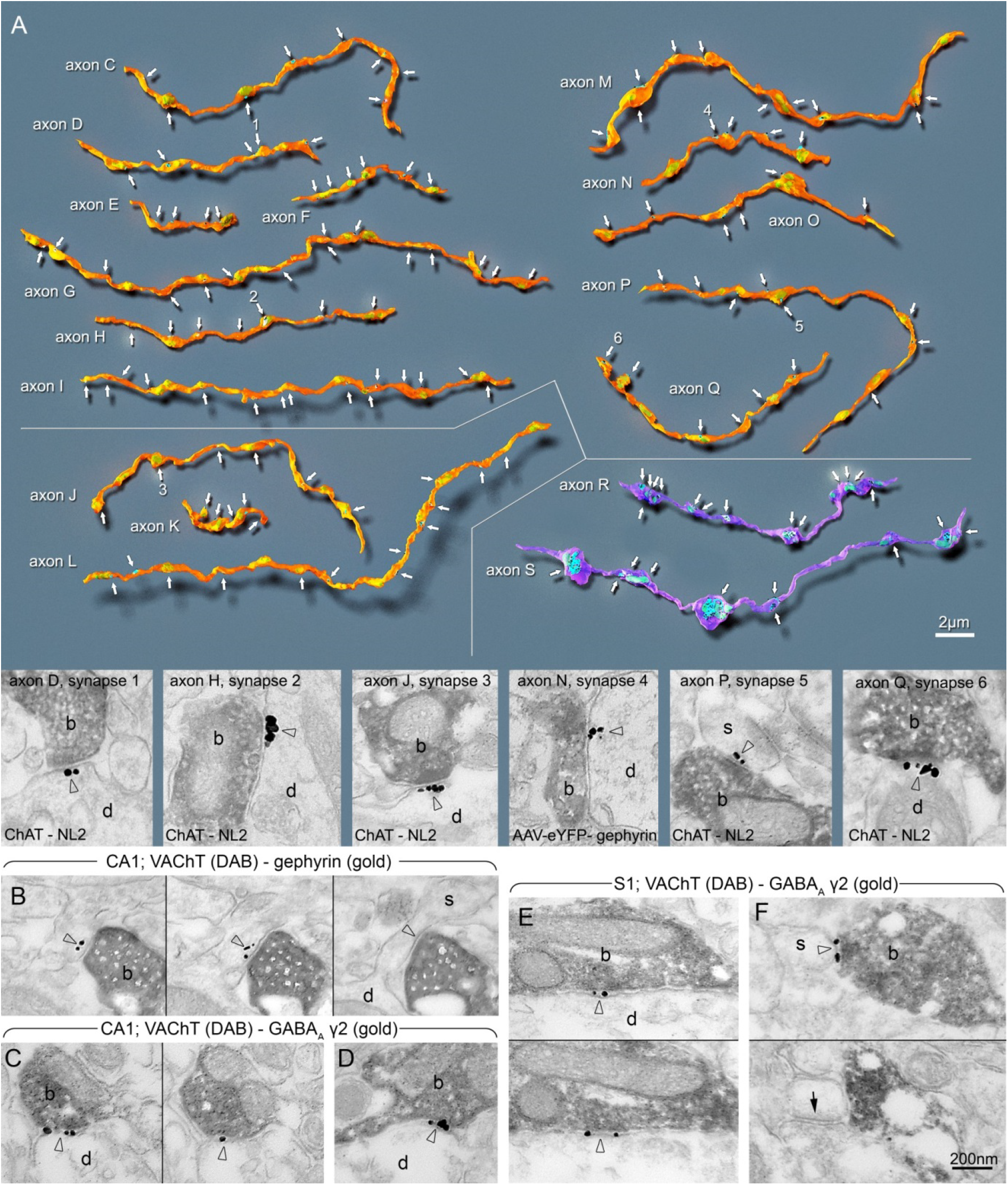
Cholinergic axons establish synaptic contacts just as frequently as other GABAergic fibers and express GABAergic postsynaptic markers. **A:** 3D EM reconstructions of DAB-labeled ChAT-positive (axons C-L; P,Q), AAV-eYFP virus-traced septo-hippocampal (axons M-O) and CB_1_-positive (axons R, S) axonal segments from the hippocampus (str. ori: axons C-F, M-O, R; str. rad: G-I, S; str. l-m: P,Q) and layer I-III of the somatosensory cortex (axons J-L). Gold labelings of NL2 (axons C-L; P-S) or gephyrin (axons M-O) were used to recognize synapses (arrows). Notice, that the linear density of synapses along cholinergic (C-Q) and CB_1_-positive GABAergic axons (R,S) are not different (see also data in Figure 2). Electron micrographs show cholinergic terminal boutons (b) forming synapses 1-6 (arrowheads, indicated by the same numbers in the 3D-reconstructions) on dendrites (d) and a spine (s). **B-F:** Electron micrographs from combined preembedding immunogold/immunoperoxidase experiments for gephyrin or GABA_A_γ2 receptor subunit (immunogold) and vAChT (DAB: dark, homogenous reaction product) reveal the presence of gephyrin postsynaptically (B, arrowheads) and GABA_A_γ2 receptor subunit (C, D; arrowheads) in the synaptic cleft of synapses established by vAChT-positive axons in the hippocampus (CA1, str. l-m: B, str. ori: C, D). E-F: vAChT-positive (DAB) cholinergic synapses express GABA_A_γ2 receptor subunits (immunogold, arrowheads) in the neocortex S1 area (E-F). Two or three consecutive sections of the same synapses are shown in B, C, E, and F. Labeled terminals shown in the EM images innervate dendrites (d) or spines (s). Spine in F receives a type I synapse (arrow in the lower panel) from an unlabeled terminal. Scale bars are 2 μm for all reconstructions and 200 nm for all EM images.

**Supplementary Figure S2.**
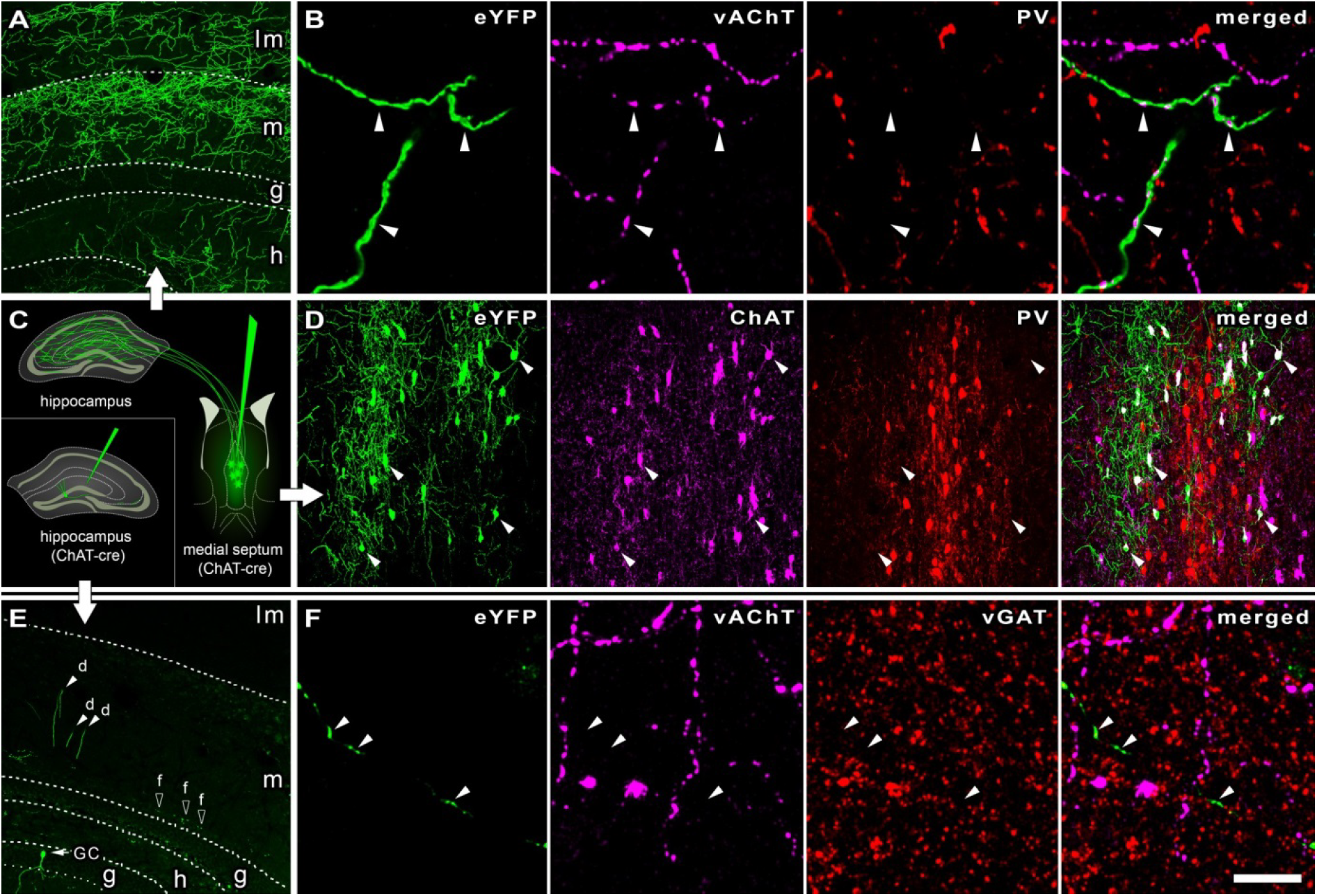
Control experiments for the labeling of septo-hippocampal cholinergic cells and fibers with Cre-dependent viral and immunolabeling techniques. **A:** A robust network of eYFP-expressing cholinergic fibers is present in the hippocampus after AAV-injection into the MS of ChAT-Cre mouse. (lm: laconosum-moleculare, m: moleculare, g: granule-cell layer, h: hilus) **B:** Confocal laser scanning microscopy images confirm that AAV-eYFP virus-traced septo-hippocampal fibers contain vAChT, but not parvalbumin (PV) in the hippocampus, demonstrating that septal GABAergic PV cells did not express Cre-dependent fluorescent protein. vAChT-labeling is localized to the terminals of the fibers. (Arrowheads mark the position of some terminals.) **C:** Schematic diagram showing the AAV-eYFP-injections into the MS or hippocampus (inset) of ChAT-Cre mice. **D:** All eYFP-expressing MS neurons were positive for ChAT, while none of them contained PV. (Arrowheads mark the position of some cell bodies.) **E:** After AAV-eYFP-injection into the hippocampus of ChAT-Cre mice, a negligible amount of cells could be detected to express eYFP. eYFP-expressing granule cell can be seen with some dendrite-segments and few scattered fibers. (d: dendrite, f: fiber, GC: granule-cell) **F:** Local eYFP-expressing fibers in the hippocampus do not contain vAChT or vGAT (Arrowheads mark some terminals, scale bar on F is 150 μm for A, D, E, and 6 μm for B and F), demonstrating the lack of cholinergic fibers originating from inside the hippocampus.

**Supplementary Figure S3.**
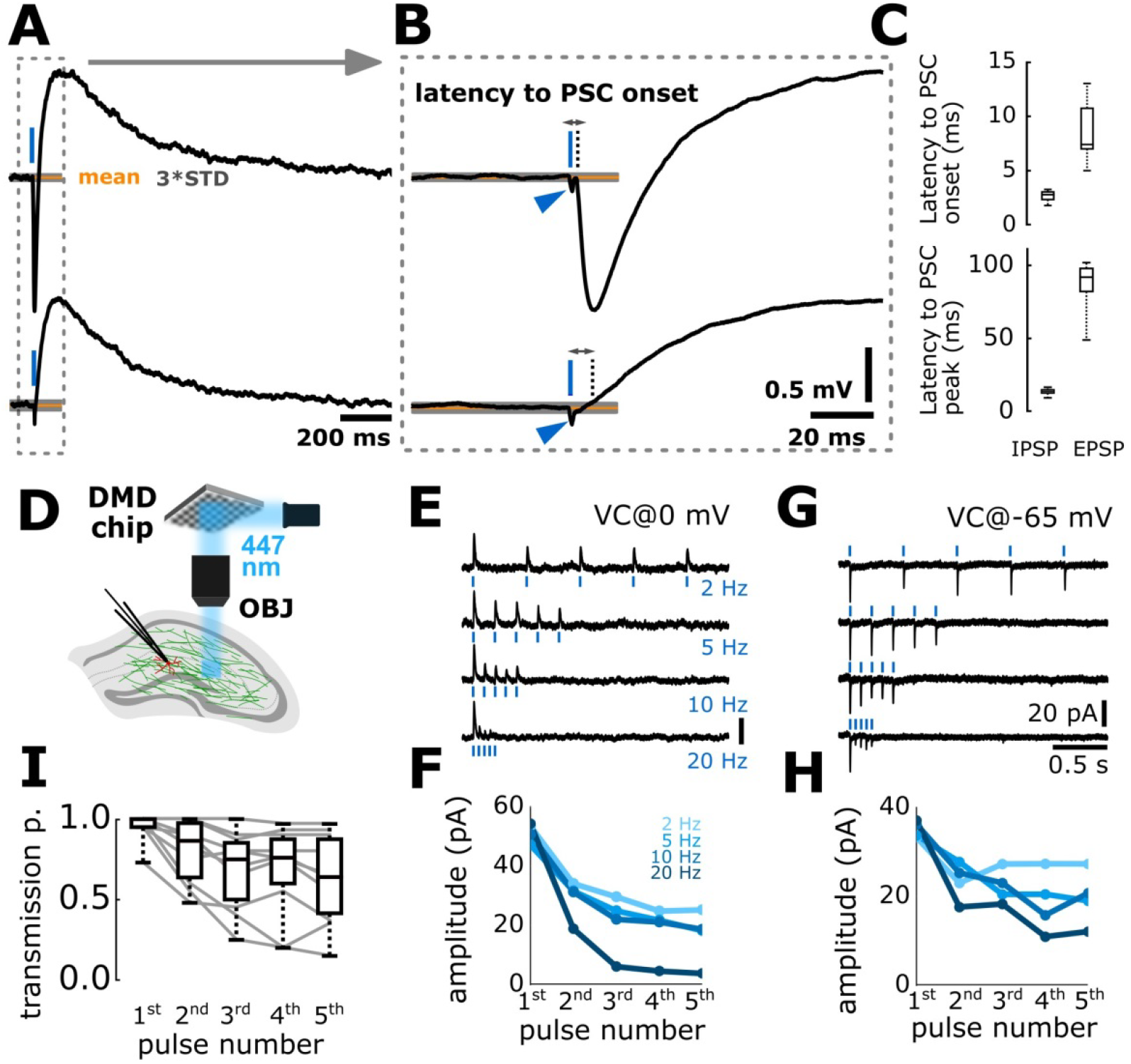
GABAergic short-term depression is a presynaptic property of cholinergic fibers. **A:** Average membrane potential response of an inhibitory neuron recorded in str. lacunosum-moleculare for cholinergic fiber stimulation (top). Latency from stimulation start to PSP onset was calculated as the time when the signal crosses 3 times standard deviation of baseline (orange line and grey shaded area represent mean and 3 times STD of the baseline). Inhibition of GABA_A_Rs (10 μM gabazine) blocks hyperpolarization (bottom) and the latency of the cholinergic response can be calculated similarly as well. **B:** Responses are magnified in time (from panel A). Blue arrowheads mark photoelectric artifacts evoked by 1 ms optical stimulus (blue bar). Note the short latency to rise (dotted line) in both the GABAergic IPSP and cholinergic EPSP. **C:** Latency from stimulus start to PSP onset (top) and PSP peak (bottom) are shown. IPSP time to onset (n = 7, in ms): 2.8 (2.2, 3.1), time to peak: 13.8 (12.7, 14.9). EPSP time to onset (n = 6, in ms): 7.4(7.0, 11.7), time to peak: 92.0 (80.5, 98.0). **D:** Channelrhodopsin 2 expression in axon terminals could change the short-term plasticity of the examined synapse by illumination driven calcium entry through the light activated channels. To exclude this possibility, with the help of a digital micro-mirror device (DMD), we have illuminated only axons running towards the measured cell, but not the axon terminals themselves. **E:** We have recorded from inhibitory neurons in str. lacunosum moleculare (n=5), and illuminated the slice far (~500 μm) from the cells. Similarly as in Figure 3, we applied 5 pulses at different frequencies (2, 5, 10 and 20 Hz). Averaged traces from one cell are shown. **F:** Mean amplitudes for different frequencies are shown from 5 cells. GABAergic currents evoked by light stimulation using the DMD device show similar short-term depression as presented on Figure 3. Amplitudes in pA (mean±std): 2 Hz: 1^st^ – 53.41(±23.66); 2^nd^ – 34.05(±17.79); 3^rd^ – 29.59(±11.54); 4^th^ – 24.80(±9.47); 5^th^ – 25.20(±11.54). 5 Hz: 1^st^ – 47.05(±21.15); 2^nd^ – 31.53(±15.58); 3^rd^ – 24.87(±12.68); 4^th^ – 21.69(±10.39); 5^th^ – 18.20(±9.58). 10 Hz: 1^st^ – 50.66(±19.43); 2^nd^ – 31.22(±13.31); 3^rd^ – 22.00(±8.16); 4^th^ – 21.10(±10.21); 5^th^ – 18.63(±4.71). 20 Hz: 1^st^ – 54.21(±18.28); 2^nd^ – 18.80(±3.95); 3^rd^ – 6.07(±1.8); 4^th^ – 4.55(±1.82); 5^th^ – 3.75(±2.11). **G:** Another factor, which can contribute to short-term depression is postsynaptic chloride loading and subsequent reduction of chloride drive, due to series of stimuli. This phenomenon can be explored with reversing chloride gradient, as demonstrated here by using a high chloride content intracellular solution. Average GABAergic responses are shown from one cell. **H:** Mean amplitudes for different frequencies are shown from 5 cells. GABAergic responses show similar short-term depression as described previously. Amplitudes in pA (mean±std): 2 Hz: 1^st^ – 33.27(±19.16); 2^nd^ – 23.02(±23.76); 3^rd^ – 27.28(±16.80); 4^th^ – 27.33(±20.33); 5^th^ – 26.31(±25.86). 5 Hz: 1^st^ – 33.27(±14.57); 2^nd^ – 27.76(±17.51); 3^rd^ – 20.46(±9.24); 4^th^ – 22.48(±10.67); 5^th^ – 10.00(±13.91). 10 Hz: 1^st^ – 36.23(±24.69); 2^nd^ – 25.29(±20.18); 3^rd^ – 23.02(±20.10); 4^th^ – 15.79(±10.36); 5^th^ – 20.76(±17.03). 20 Hz: 1^st^ – 37.09(±33.27); 2^nd^ – 17.61(±20.65); 3^rd^ – 18.29(±24.65); 4^th^ – 10.95(±10.16); 5^th^ – 12.10(±11.49). **I:** Transmission probability is shown for 10 Hz stimulation from the 10 cells presented on Figure E-H. The decrease in transmission probability support presynaptic mechanism for STD. Transmission probability: 1^st^ pulse 1(0.95, 1), 2^nd^ pulse: 0.87 (0.60, 1), 3^rd^ pulse: 0.75 (0.45, 0.87), 4^th^ pulse: 0.76 (0.55, 0.90), 5^th^ pulse: 0.64 (0.37, 0.90). Wilcoxon-sign rank test: 1^st^–2^nd^: p<0.05, 2^nd^–3^rd^: p<0.05, 3^rd^–4th and 4^th^-5^th^: not significant. In some cells, transmission probability remained stable despite the observed STD in IPSC amplitude, suggesting that multiple contacts were excited and “averaged” by optical illumination. These results support our hypothesis that GABAergic short-term depression emerges presynaptically, and not the result of channelrhodopsin-2 expression or postsynaptic chloride loading.

**Supplementary Figure S4.**
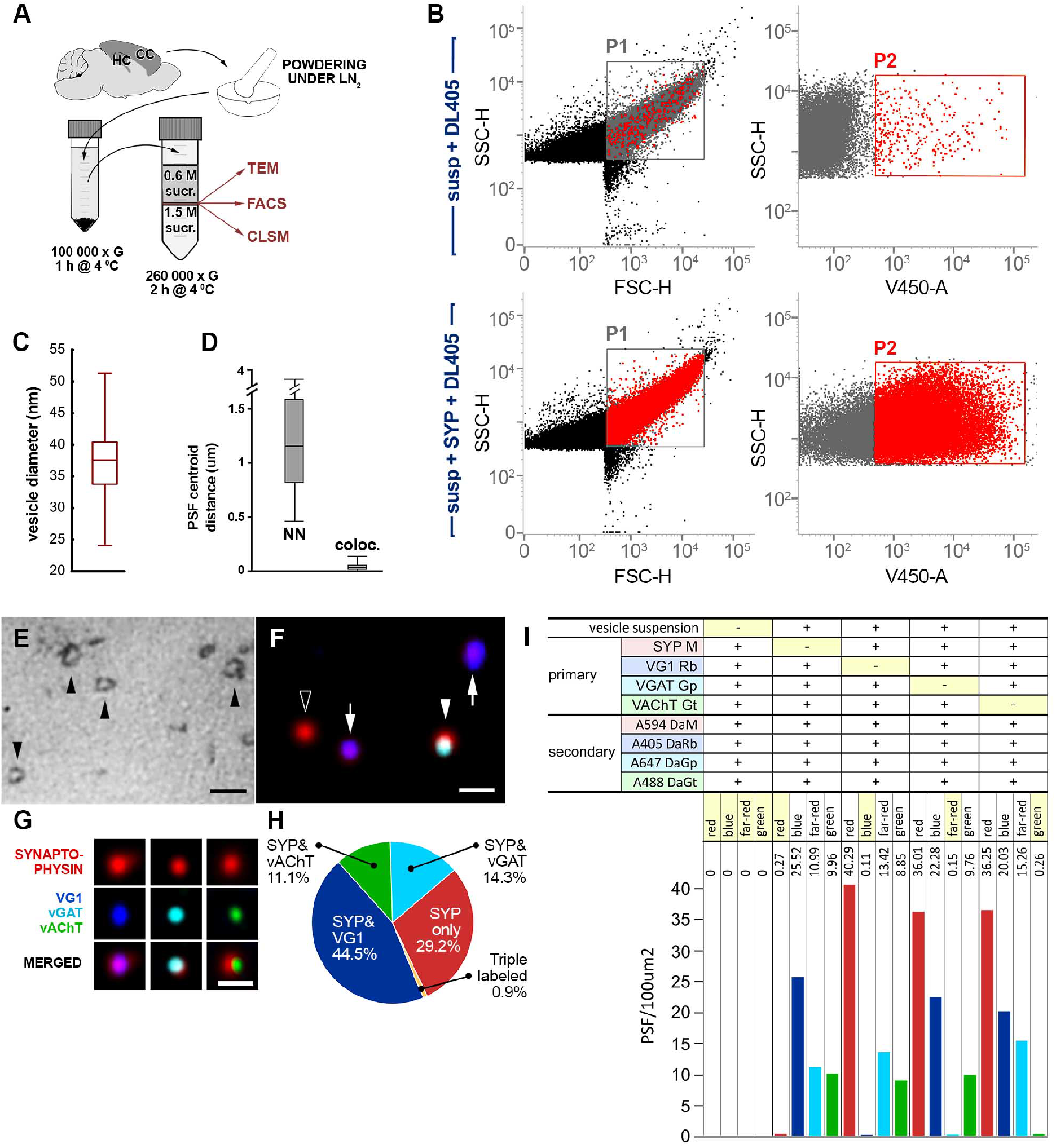
Vesicular acetylcholine and GABA transporters are localized on different vesicles. **A:** Schematic drawing depicts the main steps of synaptic vesicle isolation. Cerebral cortices (CC) and hippocampi (HC) of mice were frozen and powdered under liquid nitrogen. After homogenization, the sample was centrifuged at 100 000 G (1h, 4°C). The supernatant was laid onto a 0,6M/1,5M sucrose step gradient and centrifuged at 260 000 G (2h, 4°C). Synaptic vesicles were collected from the 0,6M/1,5M sucrose solution interface, and processed for transmission electron microscopy (TEM), flow cytometry (FACS) and confocal laser scanning microscopy (CLSM). **B:** Flow cytometric analysis of isolated synaptic vesicles from mouse hippocampal and neocortical areas show the purity of our vesicular sample preparations. Dot plot diagrams show putative vesicles investigated without (upper row) and with (lower row) synaptophysin (SYP) primary antibody labeling, followed by fluorescent secondary antibody labeling in both cases. Dot plots on the right, show strong fluorescence signal of SYP immunolabeled putative vesicles (SSC/V450), gated also on the SSC/FSC dot plots [P1], suggesting that the isolated sample is highly purified. **C:** Diameter distribution of isolated synaptic vesicles on the TEM images show typical vesicular diameters. **D:** Distribution of fluorescent point-spread functions on the CLSM images. SYP labeled vesicles were usually more than 1 μm away from each other as nearest-neighbor analysis of PSF centroids confirmed. Centroids of PSF in different channels belonging to one vesicle were never farther away from each other than 130 nm. **E:** Electron micrograph of fixed and dehydrated synaptic vesicles isolated (see A) from cerebral cortex and hippocampus. Arrows point to individual vesicles, scale bar is 100 nm. **F:** CLSM image of fixed and immunolabeled isolated synaptic vesicles. Arrows point to point spread functions (PSF) of VG1 and SYP co-labeled vesicles, arrowhead marks PSF of vGAT and SYP co-labeled vesicle, empty arrowhead shows PSF of vesicle positive only for SYP. Scale bar is 400 nm. **G:** CLSM images of individual PSFs representing synaptic vesicles from quadruple labeling experiments. Scale bar is 600 nm. **H:** Percentages of differently labeled synaptic vesicles (n=350 individual vesicles). Frequency of triple labeled vesicles were less than 1 percent, confirming that vesicular transporters for glutamate, GABA and acetylcholine are expressed by distinct vesicle populations. **I:** Control experiments of the immunolabeling method confirm the lack of non-specific staining. In the absence of vesicle suspension no PSF-s were found in the CLSM scans, and the exclusion of each primary antibody led to the selective disappearance of PSFs in the corresponding channel.

### SUPPLEMENTARY TABLES

**Supplementary Table S1.**
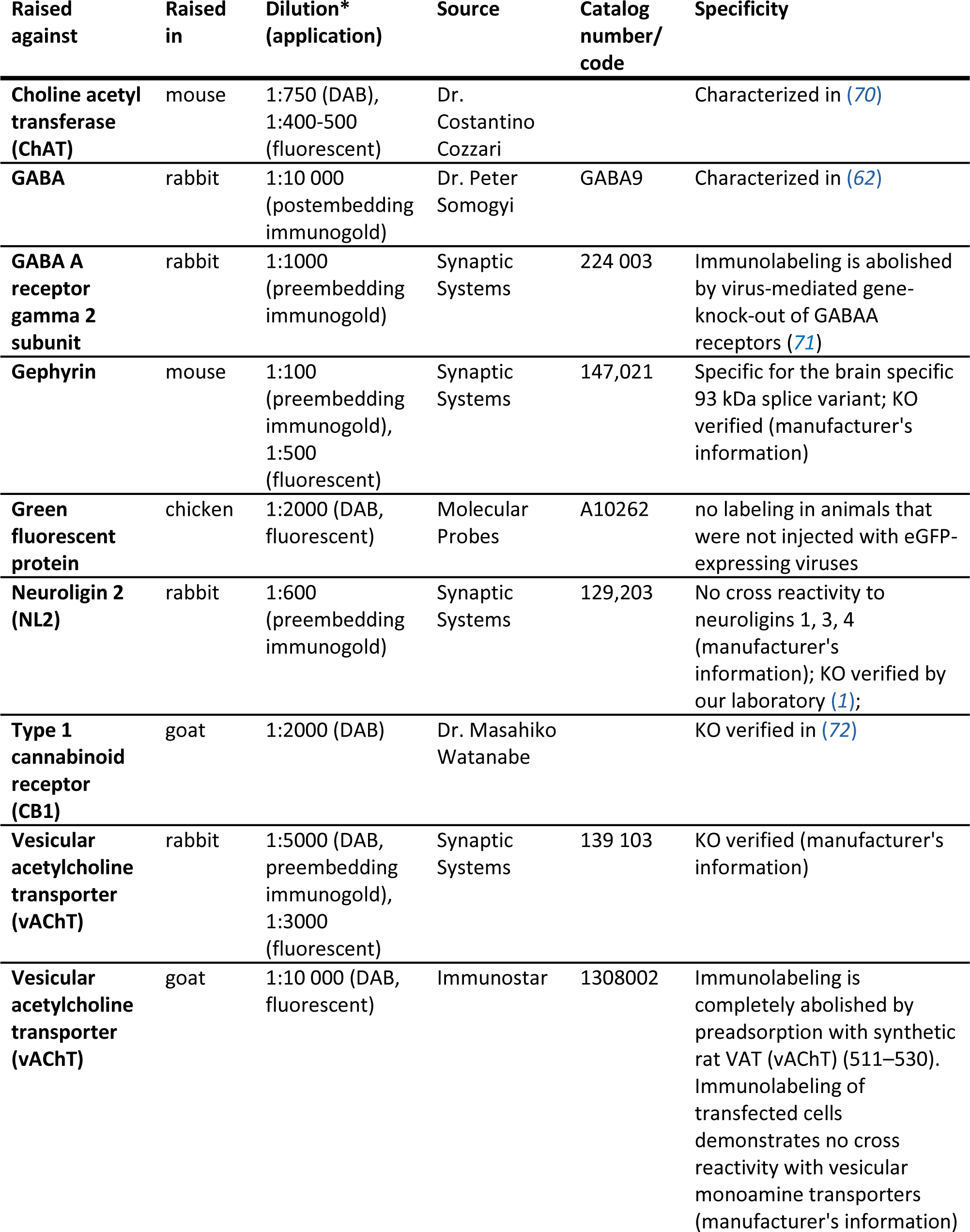
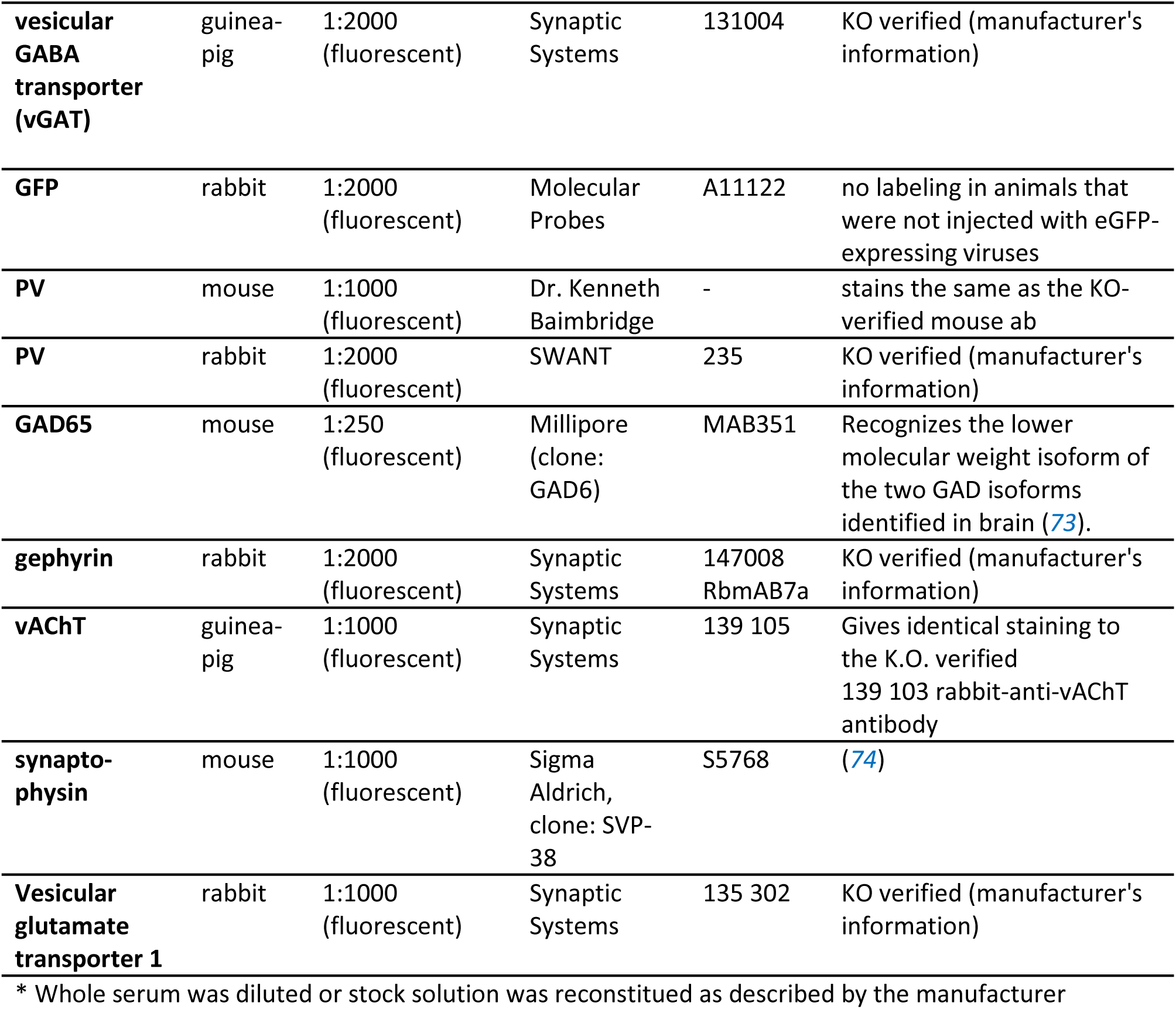
(primary antibodies)

**Supplementary Table S2.** (secondary antibodies)

| **Conjugated with** | **Raised in** | **Raised against** | **Dilution\* (procedure)** | **Source** | **Catalog number** |
| --- | --- | --- | --- | --- | --- |
| **1.4-nm nanogold** | goat | rabbit | 1:100-1:300 (preembedding immunogold) | Nanoprobes | #2004 |
| **10-nm gold** | goat | rabbit | 1:1000 (postembedding immunogold) | BBI solutions | EM.GAR10 |
| **Ultra small gold** | goat | mouse | 1:50 (preembedding immunogold) | Aurion | 100.022 |
| **Biotin-SP** | donkey | goat | 1:1000 (DAB) | JIL Inc. (Jackson Immunoresearch Laboratories, Inc.) | 705-066-147 |
| **Biotin-SP** | donkey | mouse | 1:1000 (DAB) | JIL Inc. | 715-066-151 |
| **Biotin-SP** | donkey | goat | 1:1000 (DAB) | JIL Inc. | 705-066-147 |
| **Biotin-SP** | donkey | rabbit | 1:500 (DAB) | JIL Inc. | 711-065-152 |
| **Biotin** | goat | chicken | 1:200 (DAB) | Vector Laboratories | BA-9010 |
| **DyLight 405** | donkey | mouse | 1:200 Flow cytometry | JIL Inc. | 715-475-151 |
| **DyLight 405** | donkey | rabbit | 1:1000 (confocal) | JIL Inc. | 711-475-152 |
| **Alexa 488** | goat | chicken | 1:1000 (confocal), 1:500 (STORM) | Vector Laboratories | A-11039 |
| **Alexa 488** | donkey | rabbit | 1:500 (confocal) | Invitrogen | A21206 |
| **Alexa 488** | donkey | chicken | 1:1000 (confocal) | JIL Inc. | 703-545-155 |
| **Alexa 488** | donkey | goat | 1:1000 (confocal) | JIL Inc. | 705-475-147 |
| **Cy-3** | donkey | mouse | 1:500 (confocal) | JIL Inc. | 715-165-151 |
| **Alexa 594** | donkey | rabbit | 1:500 (confocal) | Invitrogen | A21207 |
| **DyLight 549** | donkey | guinea-pig | 1:500 (confocal) | JIL Inc. | 706-505-148 |
| **Alexa 594** | donkey | mouse | 1:500 (confocal) | Life Technologies | A21203 |
| **CF 568** | donkey | guinea-pig | 1:500 (STORM) | Biotium | 20377-500uL |
| **Alexa 647** | donkey | guinea-pig | 1:500 (confocal) | JIL Inc. | 706-605-148 |
| **Cy-5** | donkey | goat | 1:500 (confocal) | JIL Inc. | 705-175-147 |
| **Alexa 647** | donkey | rabbit | 1:500 (confocal, STORM) | JIL Inc. | 711-605-152 |
| **Alexa 647** | donkey | mouse | 1:500 (confocal) | JIL Inc. | 715-605-151 |

**Supplementary Table S3.**
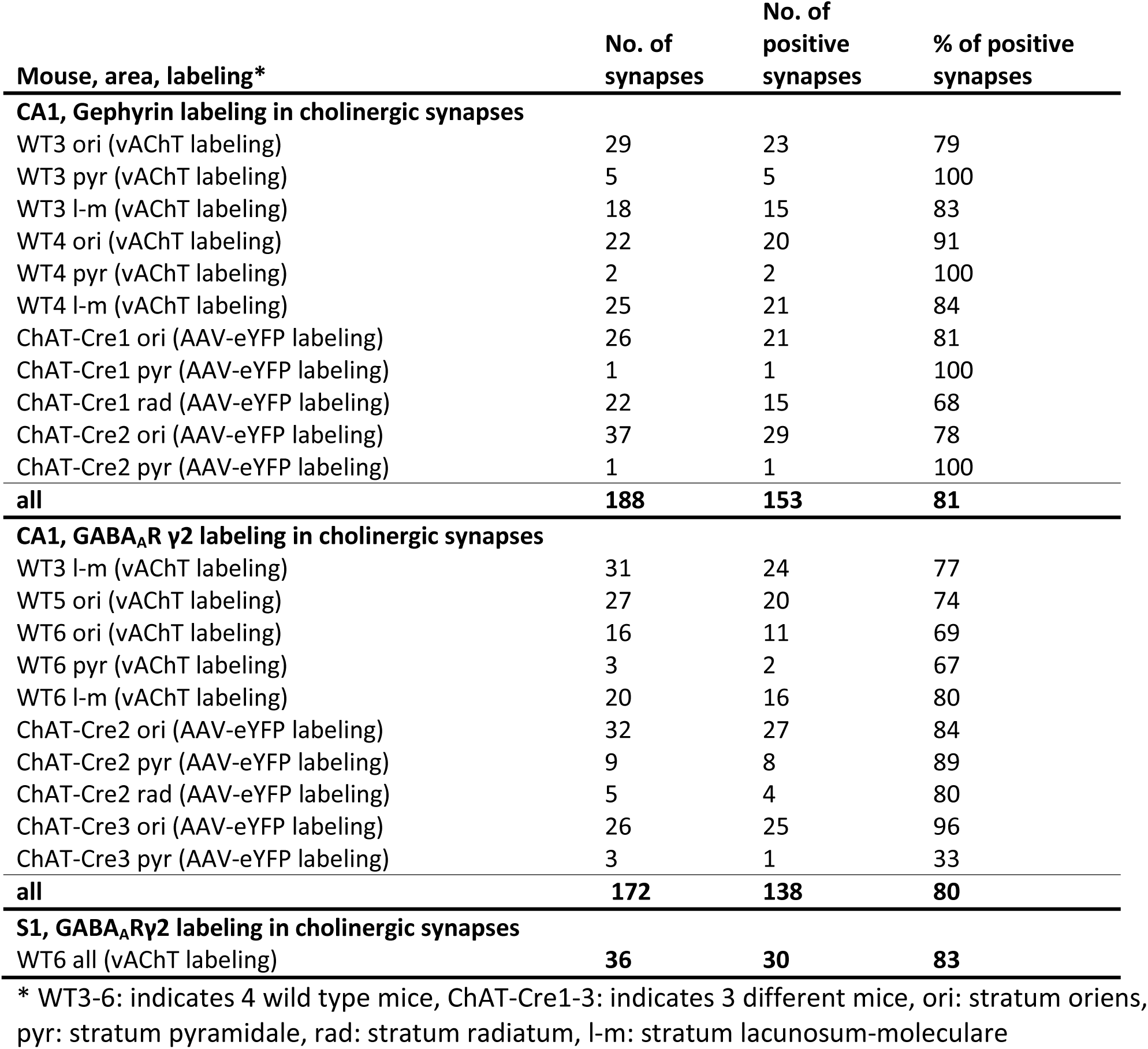
The ratio of cholinergic synapses expressing gephyrin and GABA_A_R γ2 subunit in different brain areas.

**Supplementary Table S4.** Measured parameters of reconstructed axonal segments

| Axon ID<sup>a</sup> | Labeling<br>(DAB-gold) | mouse | Length<br>(μm) | No. of mitochondria | No. of synapses | Synapse density<br>(syn./ 100 μm) | Postsynaptic targets | | | |
| --- | --- | --- | --- | --- | --- | --- | --- | --- | --- | --- |
|  |  |  |  |  |  |  | Pyramidal<br>dendrite | Interneuron<br>dendrite | Spine | Unidentified |
| A (in CA1 ori) | ChAT-NL2 | WT1 | 22.0 | 4 | 12 | 54.5 | 8 | 1 | 3 | 0 |
| C (in CA1 ori) | ChAT-NL2 | WT2 | 21.5 | 3 | 8 | 37.2 | 3 | 1 | 4 | 0 |
| D (in CA1 ori) | ChAT-NL2 | WT2 | 14.8 | 4 | 5 | 33.8 | 5 | 0 | 0 | 0 |
| E (in CA1 ori) | ChAT-NL2 | WT2 | 6.8 | 1 | 4 | 58.5 | 2 | 1 | 0 | 1 |
| F (in CA1 ori) | ChAT-NL2 | WT2 | 10.0 | 4 | 6 | 60.1 | 5 | 1 | 0 | 0 |
| B (in CA1 ori) | eYFP-gephyrin | ChAT-Cre1 | 38.1 | 11 | 15 | 39.4 | 10 | 0 | 3 | 2 |
| M (in CA1 ori) | eYFP-gephyrin | ChAT-Cre1 | 27.0 | 7 | 10 | 37.0 | 6 | 1 | 0 | 3 |
| N (in CA1 ori) | eYFP-gephyrin | ChAT-Cre2 | 12.3 | 2 | 4 | 32.6 | 4 | 0 | 0 | 0 |
| O (in CA1 ori) | eYFP-gephyrin | ChAT-Cre2 | 18.4 | 4 | 6 | 32.7 | 4 | 0 | 2 | 0 |
| **CA1 ori (cholinergic) all** |  |  | **170.9** | **40** | **70** | **41.0** | **47**<br>**(67.1%)** | **5**<br>**(7.1%)** | **12**<br>**(17.1%)** | **6**<br>**(8.6%)** |
| G (in CA1 rad) | ChAT-NL2 | WT1 | 33.1 | 12 | 15 | 45.3 | 8 | 0 | 7 | 0 |
| H (in CA1 rad) | ChAT-NL2 | WT2 | 17.2 | 4 | 7 | 40.8 | 4 | 0 | 3 | 0 |
| I (in CA1 rad) | ChAT-NL2 | WT1 | 25.4 | 9 | 15 | 59.0 | 8 | 0 | 7 | 0 |
| **CA1 rad (cholinergic) all** |  |  | **75.8** | **25** | **37** | **48.8** | **20**<br>**(54.1%)** | **0**<br>**(0%)** | **17**<br>**(45.9%)** | **0**<br>**(0%)** |
| **CA1 ori+rad (cholinergic) all** |  |  | **246.7** | **65** | **107** | **43.4** | **67**<br>**(62.6%)** | **5**<br>**(4.7%)** | **29**<br>**(27.1%)** | **6**<br>**(5.6%)** |
| P (in CA1 l-m) | ChAT-NL2 | WT2 | 26.2 | 7 | 8 | 30.5 | 2<sup>b</sup> |  | 5 | 1 |
| Q (in CA1 l-m) | ChAT-NL2 | WT2 | 17.0 | 5 | 6 | 35.3 | 2<sup>b</sup> |  | 3 | 1 |
| **CA1 l-m (cholinergic) all** |  |  | **43.2** | **12** | **14** | **32.4** | **4<sup>b</sup>**<br>**(28.6%)** |  | **8**<br>**(57.1%)** | **2**<br>**(14.3%)** |
| **CA1 cholinergic fibers in all layers** |  |  | **289.9** | **77** | **121** | **41.7** | **76<sup>b</sup>**<br>**(62.8%)** |  | **37**<br>**(30.6%)** | **8**<br>**(6.6%)** |
| J (in S1 LI) | ChAT-NL2 | WT2 | 22.0 | 7 | 6 | 27.3 | 4<sup>b</sup> |  | 2 | 0 |
| K (in S1 LII/LIII) | ChAT-NL2 | WT2 | 6.1 | 3 | 5 | 82.7 | 1<sup>b</sup> |  | 3 | 1 |
| L (in S1 LII) | ChAT-NL2 | WT2 | 32.6 | 10 | 13 | 39.9 | 8<sup>b</sup> |  | 4 | 1 |
| **S1 cholinergic fibers in all layers** |  |  | **60.6** | **20** | **24** | **39.6** | **13<sup>b</sup>**<br>**(54.2%)** |  | **9**<br>**(37.5%)** | **2**<br>**(8.3%)** |
| R (in CA1 ori) | CB<sub>1</sub>-NL2 | WT1 | 18.0 | 6 | 14 | 77.6 | 13 | 0 | 1 | 0 |
| S (in CA1 rad) | CB<sub>1</sub>-NL2 | WT1 | 29.2 | 7 | 10 | 34.2 | 8 | 0 | 0 | 2 |
| **CA1 CB<sub>1</sub>-positive fibers in all layers** |  |  | **47.2** | **13** | **24** | **50.8** | **21**<br>**(87.5%)** | **0**<br>**(0%)** | **1**<br>**(4.2%)** | **2**<br>**(8.3%)** |

<sup>a</sup>: WT3-6: indicates 4 wild type mice, ChAT-Cre1-3: indicates 3 different mice, ori: stratum oriens, pyr: stratum pyramidale, rad: stratum radiatum, l-m: stratum lacunosum-moleculare; <sup>b</sup>: We did not identify the cell types that established the dendritic shafts in CA1 lacunosum-moleculare and S1 (see Supplemental Experimental Procedures).

